# The brain of a nocturnal migratory insect, the Australian Bogong moth

**DOI:** 10.1101/810895

**Authors:** Andrea Adden, Sara Wibrand, Keram Pfeiffer, Eric Warrant, Stanley Heinze

## Abstract

Every year, millions of Australian Bogong moths (*Agrotis infusa*) complete an astonishing journey: in spring, they migrate over 1000 km from their breeding grounds to the alpine regions of the Snowy Mountains, where they endure the hot summer in the cool climate of alpine caves. In autumn, the moths return to their breeding grounds, where they mate, lay eggs and die. These moths can use visual cues in combination with the geomagnetic field to guide their flight, but how these cues are processed and integrated in the brain to drive migratory behavior is unknown. To generate an access point for functional studies, we provide a detailed description of the Bogong moth’s brain. Based on immunohistochemical stainings against synapsin and serotonin (5HT), we describe the overall layout as well as the fine structure of all major neuropils, including the regions that have previously been implicated in compass-based navigation. The resulting average brain atlas consists of 3D reconstructions of 25 separate neuropils, comprising the most detailed account of a moth brain to date. Our results show that the Bogong moth brain follows the typical lepidopteran ground pattern, with no major specializations that can be attributed to their spectacular migratory lifestyle. These findings suggest that migratory behavior does not require widespread modifications of brain structure, but might be achievable via small adjustments of neural circuitry in key brain areas. Locating these subtle changes will be a challenging task for the future, for which our study provides an essential anatomical framework.

## Introduction

One of the central questions in neuroscience is how animal behaviors result from neural computations. From the sensory input that elicits a behavior to the motor circuits that execute it – the neural processing is complex and, as yet, we only understand small pieces of this multi-level puzzle. To delve deeper into this question, we need an animal model that shows a robust behavior and whose brain is accessible enough to allow for detailed physiological and genetic studies. In terms of accessibility, insects have long been reliable model organisms (Clarac and Pearlstein, 2007), as they have comparatively small brains that control a broad and remarkably complex behavioral repertoire. In this study, we focus on one particular species: the Bogong moth *Agrotis infusa*.

The Bogong moth is one of the most iconic Australian insect species, well known for its spectacular long-distance migrations, carried out during the night (Warrant et al., 2016). Each spring, Bogong moths escape their hot breeding grounds via a 1000 km long journey to the Australian Alps, where they enter a state of dormancy (“aestivation”) inside cool, high-altitude caves (Common, 1954; Warrant et al., 2016). In early autumn, the moths return to their breeding grounds, mate, lay eggs and die. These seasonal migrations are similar to the migrations of the day-active Monarch butterfly in North America (Reppert et al., 2016), but unlike this species, which completes the migratory cycle over several generations, every Bogong moth completes the entire return journey (Common, 1954).

Crucially, if a moth loses the migratory heading during its long-distance flight, it will miss either its aestivation site (spring) or the breeding grounds (autumn). In either case it will likely not survive the journey and thus fail to reproduce. Therefore, as in all migratory animals, significant selective pressure maintains the stability and accuracy of the migratory behavior in the Bogong moth. To successfully migrate, the moths’ brains need to solve two consecutive problems: first, they must maintain a stable course across many hundreds of kilometers of unfamiliar landscape, and second, they must abort this long-range flight and search for a suitable aestivation site. None of the two processes can rely on learning, as each moth only lives through one migratory cycle. While it was found that migratory Bogong moths can use the geomagnetic field in combination with visual cues to control their flight heading (Dreyer et al., 2018), it is likely that they use olfactory cues to locate their specific aestivation caves (Warrant et al., 2016). Both the visual system and the olfactory system of the brain therefore could house adaptations that reflect migratory lifestyle. Changes in the volume and the internal structure of primary and secondary sensory brain regions that correlate with a species’ ecology have been found in hawkmoths (Stöckl et al., 2016) and social hymenopterans (Molina and O’Donnell, 2008; Amador-Vargas et al., 2015). This suggests that migratory behavior, with its high demand on reliable sensory coding during long distance flight and the need to unambiguously identify the migration target site, could also be visible as structural adaptations on the level of brain regions.

During migration, the moths’ brains have to carefully control a complex series of navigational decisions, which must be based on hardwired premotor control circuits. The central decision making centers of the brain, the central complex (CX) and the lateral complex (LX), are thus additional brain regions that could house migration-specific circuits (Honkanen et al., 2019). Although initial data from the Bogong moth have not revealed major differences in the structure of these regions when compared to a related species with no pronounced migratory behavior (*Agrotis segetum*), small volumetric changes in specific sub-regions have indeed been identified (de Vries et al. 2017). To date, these are the only indications of neural specialization potentially due to migratory lifestyle in insects. The modest nature of the observed changes in central regions suggest that most adaptations underlying the implementation of navigational strategies might be on the level of neural circuits, not neuropils (de Vries et al, 2017).

However, to carry out detailed comparative studies on the neural circuits controlling the Bogong moths’ migratory ability, and to provide the foundation for quantitative investigations of neural investments in this species, we require a detailed qualitative and quantitative description of its brain. Such a brain map will allow comparison to the brain map of the Monarch butterfly, a diurnal migratory species (Heinze et al., 2012) as well as to other, non-migratory species. It will therefore provide an access point for understanding the neural basis of long-distance, nocturnal migration. As the overall behavioral pattern of the Bogong moth migration is both highly reproducible as well as complex, a thorough understanding of its neural underpinnings will generate a simple model system that might also shed light on the sensory-motor transformations underlying other complex behaviors, including those in larger brains.

To provide the anatomical basis for those future studies, we have carried out a combination of immunohistochemical stainings and 3D reconstructions of Bogong moth brains and produced a comprehensive description of all major regions in the brain of this species, similar to those published for the Monarch butterfly (Heinze and Reppert, 2012), the dung beetle (Immonen et al., 2017), the fruit fly (Jenett et al., 2012; Ito et al., 2014), and the locust (von Hadeln et al., 2018). To account for inter-individual variation, we additionally generated a standardized brain atlas, which robustly describes the average shape of the male Bogong moth brain. Taken together, our detailed descriptions of all neuropils and the standardized brain atlas lay the groundwork for future anatomical, physiological and genetic studies of the neural basis of Bogong moth migration and navigation.

## Materials and Methods

### Animals

Aestivating Bogong moths (*Agrotis infusa*) were collected in a cave on South Ramshead (Kosciuzko National Park, New South Wales, Australia) and transported to Sweden in cooled plastic containers. In the lab, the aestivating moths were kept in a temperature-controlled incubator (I-30 VL, Percival/CLF Plant Climatics, Wertingen, Germany) in cave-like conditions (16 hours dim light at 10 °C, 8 hours dark at 6 °C). The moths had free access to food solution (2% sugar, 2% honey, 0.2% ascorbic acid in water).

In the experiments described below, female and male moths were used. At the time of the experiment, Bogong moths were several months old, as the adults survive up to 8-9 months in the wild if they estivate during the summer (Warrant et al., 2016).

### Antibodies

In the following histology protocols, we used antibodies against synapsin and serotonin (5-HT). Both antibodies are described in detail in Table 1. The anti-synapsin antibody was kindly provided by Dr. E. Buchner and Dr. C. Wegener (Würzburg University, Germany; Cat# SYNORF1 (*Drosophila* synapsin I isoform), RRID: AB_2315426; (Klagges et al., 1996)). For stainings against serotonin we used the 5-HT rabbit antibody from Immunostar (Hudson, WI, USA; Cat# 20080, Lot# 051007). Please note that anti-serotonin labeling was exclusively used to highlight structural detail of brain regions rather than to discuss the expression patterns of serotonin. The secondary antibodies were GAM-Cy5 (goat anti mouse conjugated to Cy5; Jackson Immunoresearch Laboratories Inc., West Grove, PA, USA; Cat# 115-175-146, Lot# 108262) and GAR-488 (goat anti rabbit conjugated to Alexa Fluor 488; Invitrogen, Eugene, OR, USA; Cat# A-11008, Lot# 57099A). Normal goat serum (NGS; Jackson Immunoresearch, West Grove, PA, USA; Cat# 005-000-121, Lot# 126560) was used for blocking non-specific antibody binding sites.

**TABLE 1.**
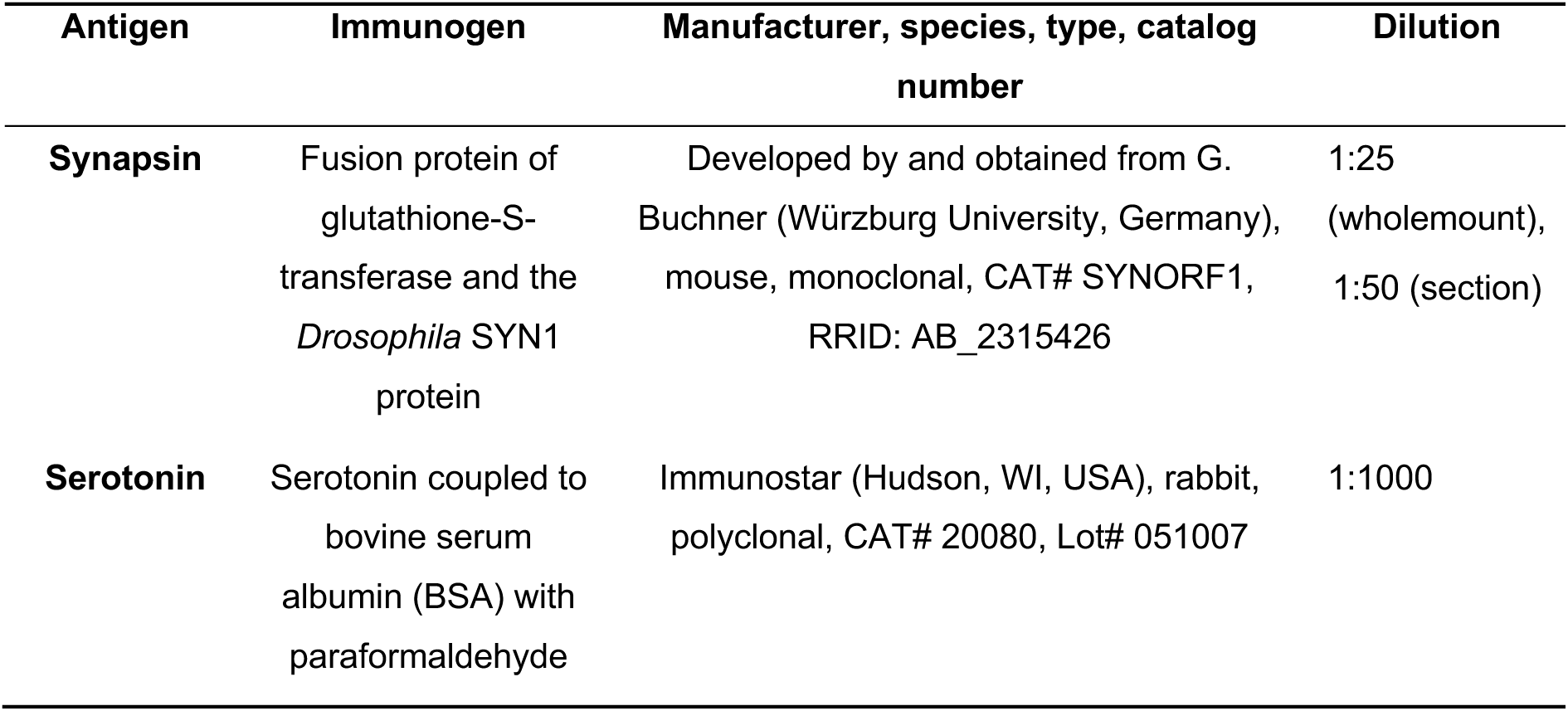
Primary antibodies.

| Antigen | Immunogen | Manufacturer, species, type, catalog number | Dilution |
| --- | --- | --- | --- |
| **Synapsin** | Fusion protein of glutathione-S-transferase and the *Drosophila* SYN1 protein | Developed by and obtained from G. Buchner (Würzburg University, Germany), mouse, monoclonal, CAT# SYNORF1, RRID: AB_2315426 | 1:25<br>(wholemound),<br>1:50 (section) |
| **Serotonin** | Serotonin coupled to bovine serum albumin (BSA) with paraformaldehyde | Immunostar (Hudson, WI, USA), rabbit, polyclonal, CAT# 20080, Lot# 051007 | 1:1000 |

### Immunohistochemistry protocols

The whole-mount staining protocol was adapted from the protocols described in Ott (2008), Heinze and Reppert (2012) and Stöckl and Heinze (2015). Moth heads were mounted in a wax-filled petri dish and dissected to expose the brain. Fresh zinc-formaldehyde fixative (18.4 mM ZnCl_2_, 135 mM NaCl, 35 mM sucrose, 1% PFA; (Ott, 2008)) was immediately applied to the brain, which was then dissected out of the head capsule and cleaned of trachea and fat body. Subsequently, the retinae were removed from the isolated brains. Brains were left to fix for 20 hours at 4°C and were then washed 8×20 minutes in HEPES-buffered saline (HBS: 150 mM NaCl, 5 mM KCl, 5 mM CaCl_2_, 25 mM sucrose, 10 mM HEPES; (Ott, 2008)). Brains were bleached in 10% H_2_O_2_ in Tris-buffered saline (Tris-HCl) for 8 hours (Stöckl & Heinze, 2015). For some preparations, the bleaching step was omitted. Subsequently, brains were washed in Tris-HCl (3×10 minutes), permeabilized in a fresh mixture of dimethyl sulfoxide (DMSO) and methanol (20:80, 70 minutes) and washed again in Tris-HCl (3×10 minutes). After pre-incubating the brains in 5% normal goat serum (NGS) in 0.01 M phosphate-buffered saline (PBS) with 0.3% TritonX-100 (PBS-Tx) over night at 4°C, brains were transferred to the primary antibody solution (anti-synapsin 1:25, 1% NGS in PBS-Tx) and left to incubate at 4°C for 6 days. Brains were washed 8×20 minutes in PBS-Tx before being transferred to the secondary antibody solution (1:300 GAM-Cy5, 1% NGS in PBS-Tx) and left to incubate at 4°C for 5 days. Before mounting, brains were washed in PBS-Tx (6×15 minutes) and PBS (2×15 minutes) and dehydrated in an increasing ethanol series (50%, 70%, 90%, 96%, 2×100%, 15 minutes each). For clearing, brains were transferred to a fresh mixture of ethanol and methylsalicylate (1:1) and the ethanol was allowed to evaporate. After 15 minutes, the mixture was replaced by pure methylsalicylate, in which the brains were left to clear for up to 75 minutes. Finally, the brains were embedded in Permount (Electron Microscopy Science, Hartfield, PA, USA) between two coverslips, using plastic spacers (Zweckform No. 3510, Germany) to prevent squeezing.

For brain sections, we used a protocol adapted from the sectioning protocol described in (Heinze and Reppert, 2012). Brains were dissected in moth ringer solution (150 mM NaCl, 3 mM KCl, 10 mM TES, 25 mM sucrose, 3 mM CaCl_2_; based on (King et al., 2000)), immediately transferred to Zamboni’s fixative (4% PFA, 7.5% picric acid in 0.1 M phosphate buffer) and left to fix over night at 4°C. The brains were then washed in PBS (3×10 minutes), embedded in albumin-gelatin (4.8% gelatin and 12% ovalbumin in demineralized water) and post-fixed in 4% formaldehyde solution (in 0.1 M PBS). A vibratome (Leica VT1000 S, Leica Biosystems, Nussloch, Germany) was used to cut the brains into 40 *µ*m thick sections, which were then washed in 0.1 M PBS and pre-incubated in 5% NGS in PBS-Tx (1 hour) before being transferred to the primary antibody solution (1:50 anti-synapsin, 1:1000 anti-5HT, 1% NGS in PBS-Tx, incubate over night at room temperature). The sections were rinsed in PBS-Tx (8×10 minutes) and transferred to the secondary antibody solution (1:300 GAM-Cy5, 1:300 GAR-488, 1% NGS in PBS-Tx, incubated over night at room temperature). Before mounting, the sections were washed in PBS-Tx (3×10 minutes) and then mounted on chromalaun/gelatine-coated glass slides and left to dry for at least 5 hours. Once dry, the sections were dehydrated in an increasing ethanol series (demineralized water (5 minutes); 50%, 70%, 90%, 96%, 2×100% ethanol (3 minutes each)), cleared in xylene (2×5 minutes) and embedded in Entellan (EMS, Hatfield, PA).

For high resolution image stacks of immunolabeled brains, thick sections were prepared. In the current study these were obtained from neurobiotin injected brains that had previously been imaged at low resolution. These brains were incubated in xylene for several hours to remove the Permount mounting medium. Then the brains were rehydrated in a series of decreasing ethanol concentrations (15 minutes each, reverse order from above). After washing the brains in 0.01 M PBS (3×15 minutes), they were embedded in albumin-gelatin and post-fixed as described above. Sections of 140 *µ*m thickness were obtained with a vibrating blade microtome. These thick sections were subjected to immunohistochemistry against synapsin and 5TH in a similar manner to whole-mount preparations. Free floating sections were rinsed in PBS (3×15 minutes), pre-incubated with 5% NGS in PBT (5 hours) and subjected to primary antibody incubation (2-3 days at 4°C; concentrations as above). After rinsing with PBT (6×15 minutes), the secondary antibody incubation was performed (1-2 days at 4°C; concentrations as above). The sections were then rinsed (4×15 minutes PBT and 2×15 minutes PBS), before being dehydrated in an increasing series of ethanol concentrations (10 minutes each step, concentrations as above). Finally, the sections were cleared in Methylsalicylate and mounted in Permount between two coverslips, separated by spacers.

### Classic histology and dye injections

For Azur II-methylene blue staining, brains were fixed over night at 4°C (in 2% paraformaldehyde, 2.5% glutaraldehyde, 2mM CaCl_2_ in 0.1M sodium cacodylic buffer). This was followed by rinsing in 0.1M sodium cacodylic buffer, dehydration in an increasing ethanol series (70% 2×10 minutes, 96% 2×10 minutes, 100% 2×15 minutes) and infiltration in an acetone/Epon plastic series (acetone 2×20 minutes, 2:1 acetone/Epon 1 hour, 1:1 acetone/Epon over night, pure Epon 6 hours). The samples were embedded in fresh Epon and polymerised at 60°C for 48 hours. Samples were then sectioned in 1 *µ*m sections on a Leica Ultracut UCT ultramicrotome using a diamond histoknife. The sections were mounted on chromalum-gelatine coated microscope slides and stained with one drop of Azur II-methylene blue.

Single neurons were injected with 4% neurobiotin solution (in 1M KCl) using a sharp micropipette during intracellular recordings (for details see e.g. (Stone et al., 2017)). Mass dye injections were performed with glass micropipettes whose tips had been broken off with a pair of tweezers. The tips of the resulting blunt micropipettes were coated in petroleum jelly and dipped into neurobiotin powder. Injections were then carried out by manually inserting the micropipette into the region of interest (antennal lobe or optic lobe). For both types of injections, the brains were dissected out of the head capsule in moth ringer solution and subsequently fixed in neurobiotin fixative (4% paraformaldehyde, 0.25% glutaraldehyde, 2% saturated picric acid in 0.1M phosphate buffer) over night at 4°C. Brains were then washed 4×15 minutes in 0.1M PBS and left to incubate in streptavidin-Cy3 or streptavidin-Cy5 (1:1000 in PBT) at 4°C for three days. After washing 6×20 minutes in PBT and 2×20 minutes in PBS, the protocol then followed the wholemount staining protocol for dehydration, clearing and mounting.

### Confocal imaging

Images were taken with a confocal laser scanning microscope (LSM 510 Meta, Zeiss, Jena, Germany, or Leica TCS SP8, Leica Microsystems, Wetzlar, Germany).

For 3D reconstructions of anti-synapsin labelled preparations, whole-mount brains were imaged with the Zeiss LSM 510 Meta, using the 633nm HeNe laser and a 10x objective (C-Apochromat 10x/0.45W, Zeiss) or, for high resolution scans, using a 25x objective (LD LCI Plan-Apochromat 25x/0.8 DIC Imm Corr, Zeiss). Scans from anterior and posterior were later aligned and merged in the 3D reconstruction software Amira 5.3.3 (FEI, Hillsboro, OR, USA). Scans of whole-mount preparations were acquired as stacks of 1 *µ*m thick optical sections at 1024×1024 resolution. The detector range was set to 650-700 nm and the pinhole was set to 1 airy unit. Detector gain and offset as well as laser power were adjusted for each section in order to optimally expose the image.

High-resolution scans of neurobiotin-labelled neurons were obtained with a 63x glycerol immersion objective (HC PL APO CS2 63x/1.30 GLYC) on a Leica TCS SP8 microscope. Sectioned brains were scanned using the 25x objective or, for detailed images, using the 40x objective (Plan-Neofluar 40x/1.3 Oil DIC, Zeiss), using the 633nm HeNe and the 488nm Ar lasers. Using the hybrid detector (HyD^TM^) of the Leica TCS SP8 microscope, high resolution scans of sections were obtained with the 20x objective (HC PL APO 20x/0.75 IMM CORR CS2) or the 63x objective (HC PL APO 63x/1.40 Oil CS2). Section images were acquired as stacks of 1*µ*m thick optical sections at 1024×1024 resolution. The detector range was set to 650-700 nm for Cy5 and 500-600 nm for Alexa488 labelled sections. The two channels were always scanned sequentially. The pinhole was set to 1 airy unit.

Scans were aligned and merged using the stitching tool in the ImageJ implementation FIJI (general public license, downloadable from http://fiji.sc (Schindelin et al., 2012); stitching algorithm based on (Preibisch et al., 2009)). Images were generally optimized for brightness and contrast. Some images were denoised using the 3D hybrid median filter algorithm implemented in FIJI (https://imagej.nih.gov/ij/plugins/hybrid3dmedian.html). All confocal images shown in this paper are single optical slices (unless stated otherwise). The terms dorsal, ventral, anterior and posterior refer to the body axis of the animal.

### Three-dimensional reconstructions

The neuropil reconstructions shown in this paper were done in the 3D reconstruction software Amira 5.3.3, and are based on anti-synapsin labelled whole-mount brains. Image stacks were down-sampled to a voxel size of 2×2×2 *µ*m. Each voxel was then assigned to an individual brain area using the segmentation editor of Amira, and the full structure of reconstructed neuropils was interpolated using the “wrap” function. Neuropil volumes were extracted using Amira’s “MaterialStatistics” function. We generated a polygonal surface model to visualize the neuropils.

### Standardization

To generate the standard confocal image stack, we used the computational morphometry toolkit (CMTK 3.2.3) and the iterative shape averaging (ISA) protocol to create an average image stack from 10 individual image stacks (Rohlfing et al., 2001). Calculations were carried out at the University of Marburg on the high-performance Linux cluster MaRC2, using a 64 core (4x AMD Opteron 6276 Interlagos) node with 4 GB of RAM per core. Of the 10 individual brains, we chose the one that was the most representative of the population in terms of volume and shape as the reference for standardization. All other brains were first affinely registered to the reference brain, to compensate for differences in size, position and rotation. After affine registration, an average image stack was computed from all 10 brains, which served as a template for the subsequent elastic registration processes. Elastic registration applies local transformations to each brain, thereby optimizing the similarity between images. The resulting images were used to compute a new average image stack, which was the template for a second elastic registration. This was repeated five times. The final average image stack was accepted as the standard image stack, and the registration parameters that were applied to each brain were then applied to the corresponding neuropil reconstructions. Finally, we used the shape-based averaging method to obtain the standard surface reconstruction (Rohlfing and Maurer, 2007). A detailed description of the method can be found in (de Vries et al., 2017). Since the lamina was only intact in three specimens, the average lamina was generated separately and later added to the standard brain. Furthermore, the unstructured protocerebrum shown in this paper was reconstructed after standardization from the standard confocal image stack. During standardization of the neuropil surfaces, the anterior and posterior parts of the right Y-tract became separated, which was fixed manually after standardization by re-reconstructing this part of the neuropil from the average image stack.

### Data availability statement

The reconstructed male standard brain and the individual female brains, as well as the standard confocal image stack, are freely available on insectbraindb.org, species handle https://hdl.handle.net/20.500.12158/SIN-0000002.1. All other raw data are available upon request.

### Results

The aim of the anatomical data presented in this study is to provide a reliable framework for future anatomical and functional work in the Bogong moth. Beyond describing each neuropil, our data also allows the direct comparison of anatomical data from many individual brains, for instance data resulting from intracellular recordings combined with dye injections. To this end we have generated a standardized atlas of the male Bogong moth brain (Figure 1). Data from individual brains can be reliably registered into this standard brain, as inter-individual differences in size and shape have been averaged. This average brain additionally provides a reference data-set of representative neuropil volumes, that is, a basis for volumetric comparisons with other insects (Figure 2). The male standard brain is the average of 10 individual brains, generated by the iterative shape averaging (ISA) protocol, using the CMTK toolkit (Rohlfing et al., 2001). As the lamina of the optic lobe was only intact in three individuals, we averaged those separately and combined the resulting average with the standard brain. The undefined neuropils of the central brain were reconstructed as a surface model based on the average image stack rather than in each individual brain.

**FIGURE 1.**
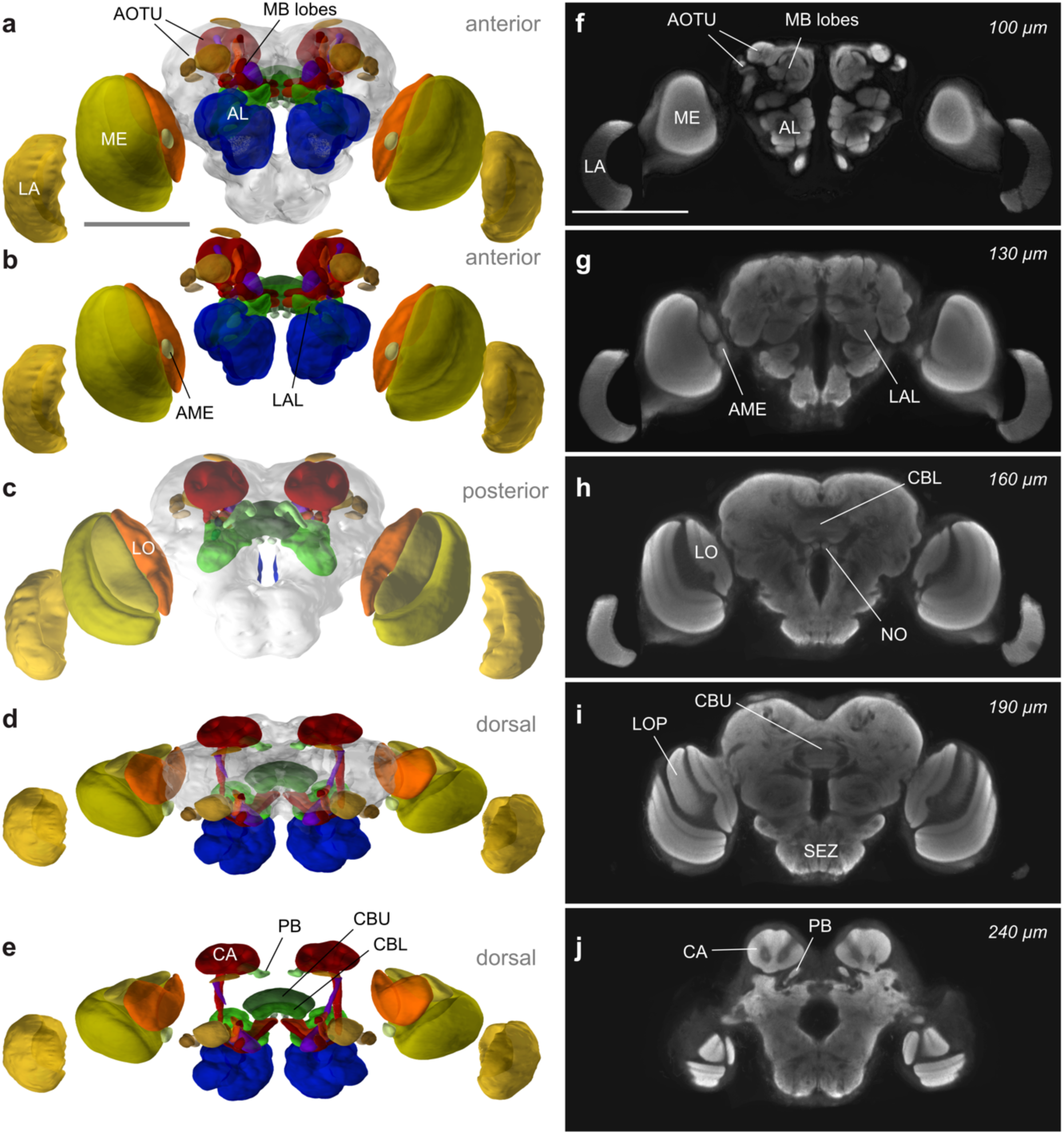
The Bogong moth standard brain. (a) Anterior view of the male standard brain reconstruction. Here and in all following figures, the colour code is as follows: yellow – visual input areas (optic lobes and ocellar neuropils), brown – anterior optic tubercle, red – mushroom bodies, green – central complex and lateral complex, blue – antennal lobes, grey – unstructured protocerebrum. (b) Anterior view without the unstructured protocerebrum, to show the arrangement of neuropils within the central brain. (c) Posterior view of the brain, with the lobula plate, ocellar neuropils, mushroom body calyces and protocerebral bridge visible. (d, e) Dorsal view of the brain with and without the unstructured protocerebrum. (f-j) Optical sections through the standard brain progressing from anterior to posterior. Note that the lamina of the standard brain was calculated separately from three available specimens and added to the standard brain later. Scale bars = 500 *µ*m.

**FIGURE 2.**
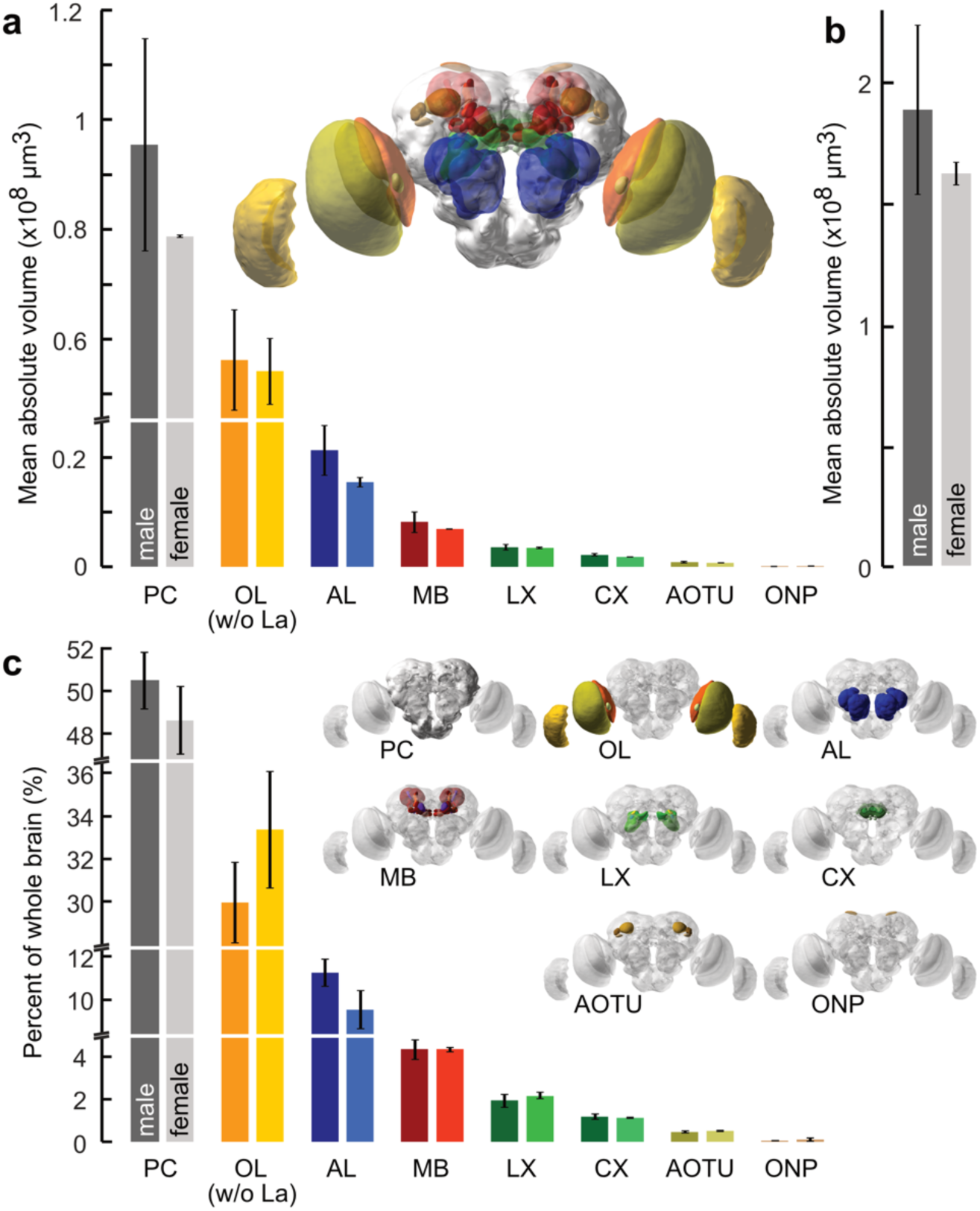
Volumetric analysis of the Bogong moth brain. (a) Mean absolute volumes of the neuropils within the male and female Bogong moth brain. (b) Mean absolute volumes of the complete male and female brain, excluding the lamina. (c) Comparison of the volume contributions of each neuropil to the whole brain, for 6 male and 2 female Bogong moths. Note that the optic lobes exclude the lamina, which was only intact in four samples (three males and one female).

In total, we have reconstructed 25 individual neuropils (23 paired and 2 unpaired). These include five compartments of the optic lobes (OL), the antennal lobes (AL), three subunits of the anterior optic tubercle (AOTU), eight neuropils of the mushroom bodies (MB), four compartments of the central complex (CX), three compartments of the lateral complex (LX), and the combined mass of the remaining undefined neuropils of the central brain.

Complete volumetric analysis was carried out in six out of the ten male brains used for shape averaging and in two additional female individuals. The male Bogong moth brain was found to be slightly larger than the female, with overall volumes of 0.188 ± 0.035 mm^3^ (males) and 0.162 ± 0.005 mm^3^ (females; Table 2, Figure 2b). When normalizing to overall brain volume, we found no gross differences between the male and female brain, with the notable exception of the antennal lobe, which displayed a male-specific macroglomerular complex (MGC), which is a pronounced sexual dimorphism typical for moths (Berg et al., 1998; Stöckl et al., 2016). In relative terms, the undefined protocerebrum makes up the largest part of the brain, accounting for ca. 50% of the total volume (Table 3, Figure 2c). The second most voluminous neuropil group is the optic lobe, occupying about one third of the total brain volume. Half of the remaining 20% of brain volume is occupied by the antennal lobes. Finally, all other brain regions combined (mushroom body, lateral complex, CX, AOTU, ocellar neuropils) account for ca. 10% of total brain volume. The inter-individual variation within the population of our wild-caught Bogong moth sample was comparably high. Across the six male brains, neuropil volumes varied between 13% and 56% around the mean (Table 3), with the smallest neuropils having the largest relative variability.

**TABLE 2.**
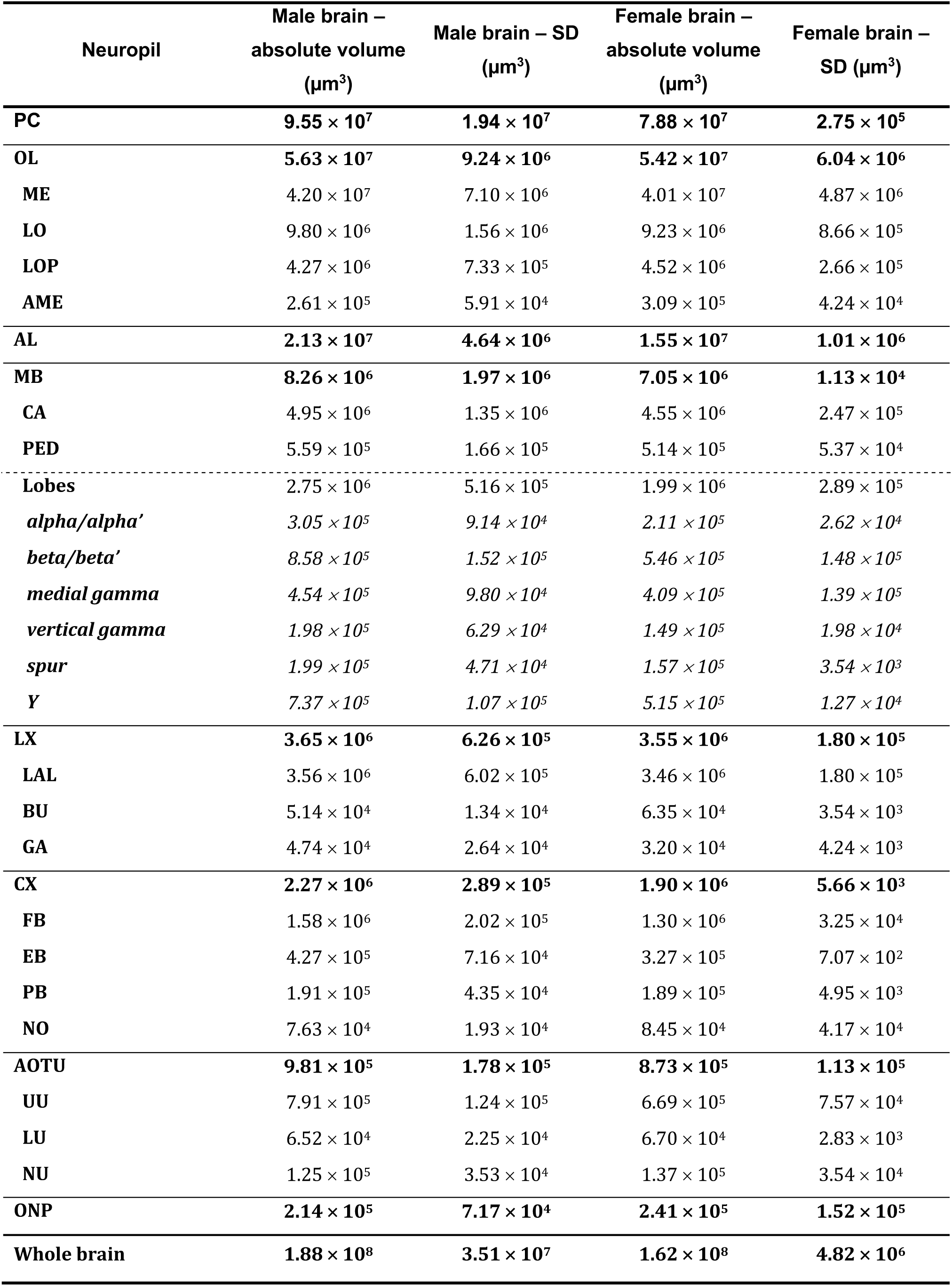
Mean absolute volumes of male and female Bogong moth brains and neuropils.

**TABLE 3.** Mean relative volumes of the neuropils of six male and two female Bogong moth brains. The normalised standard deviation (SD) is the percentage by which the volume varies around the mean (absolute SD / absolute mean x 100).

| Neuropil | Male brain –<br>relative volume<br>(%) | Male brain –<br>normalised SD<br>(%) | Female brain –<br>relative volume<br>(%) | Female brain –<br>normalised SD<br>(%) |
| --- | --- | --- | --- | --- |
| **PC** | **50.51** | **20.30** | **48.65** | **0.35** |
| **OL** | **30.01** | **16.42** | **33.40** | **11.14** |
| **ME** | 22.36 | 16.92 | 24.73 | 12.12 |
| **LO** | 5.24 | 15.91 | 5.69 | 9.39 |
| **LOP** | 2.27 | 17.17 | 2.79 | 5.88 |
| **AME** | 0.14 | 22.64 | 0.19 | 13.73 |
| **AL** | **11.28** | **21.72** | **9.55** | **6.56** |
| **MB** | **4.36** | **23.89** | **4.35** | **0.16** |
| **CA** | 2.60 | 27.39 | 2.81 | 5.44 |
| **PED** | 0.29 | 29.65 | 0.32 | 10.46 |
| **Lobes** | 1.47 | 18.76 | 1.22 | 14.57 |
| *alpha/alpha'* | 0.16 | 30.00 | 0.13 | 12.43 |
| *beta/beta'* | 0.46 | 17.75 | 0.34 | 27.09 |
| *Medial gamma* | 0.24 | 21.62 | 0.25 | 33.89 |
| *Vertical gamma* | 0.10 | 31.76 | 0.09 | 13.29 |
| *Spur* | 0.11 | 23.69 | 0.10 | 2.26 |
| *Y* | 0.40 | 14.49 | 0.32 | 2.47 |
| **LX** | **1.97** | **17.13** | **2.19** | **5.06** |
| **LAL** | 1.92 | 16.94 | 2.13 | 5.20 |
| **BU** | 0.03 | 26.00 | 0.04 | 5.57 |
| **GA** | 0.02 | 55.71 | 0.02 | 13.26 |
| **CX** | **1.22** | **12.70** | **1.17** | **0.30** |
| **FB** | 0.85 | 12.77 | 0.80 | 2.51 |
| **EB** | 0.23 | 16.75 | 0.20 | 0.22 |
| **PB** | 0.10 | 22.85 | 0.12 | 2.63 |
| **NO** | 0.04 | 25.29 | 0.05 | 49.37 |
| **AOTU** | **0.53** | **18.18** | **0.54** | **12.96** |
| **UU** | 0.42 | 15.72 | 0.41 | 11.32 |
| **LU** | 0.03 | 34.52 | 0.04 | 4.22 |
| **NU** | 0.07 | 28.13 | 0.08 | 25.81 |
| **ONP** | **0.11** | **33.51** | **0.15** | **63.21** |

In the following, all brain regions will be described in detail, starting with the primary sensory areas and moving towards more central, higher-order regions.

### Optic neuropils

The Bogong moth possesses two types of eyes: firstly, the large superposition compound eyes on either side of the head, and secondly, two ocelli, which are small lens eyes located just dorsally of the compound eyes.

Underlying the retinae of the compound eyes, the Bogong moth optic lobes contain four retinotopically organized neuropils: three large neuropils stacked from distal to proximal (lamina, medulla, lobula), and the posteriorly located lobula plate. Finally, a small fifth neuropil, the accessory medulla, is found at the anterior margin of the lobula.

The most distal neuropil is the lamina (Figure 3a, e). It consists of two distinct layers, a thick outer layer and a thin inner layer. The latter is characterized by intense synapsin immunoreactivity. This neuropil is the first processing stage of visual input and is connected to the second processing stage, the medulla, via the outer optic chiasm (Figure 3e). The medulla is the largest neuropil in the optic lobe and can be subdivided into an outer and inner medulla. Anti-synapsin and anti-5HT labelling demonstrates an intricate layering of both compartments. Overall, ten medulla layers can be distinguished in this way, of which seven are located in the outer medulla and three are in the inner medulla (Figure 3c). The serpentine layer separates the outer and inner medulla and provides an entry point for tangential fibers innervating both parts. This layer also exhibits the most intense 5HT labelling within the medulla, which otherwise is mostly present in the inner medulla, with only very sparse 5HT immunoreactive terminals present across the outer medulla. Separated by the inner optic chiasm, the lobula lies medially of the medulla and consists of three layers (Figure 3g). Unlike in hawkmoths, there is no pronounced separation into an outer and inner lobula (el Jundi et al., 2009). In line with this observation, 5HT labelling is evenly distributed across the entire neuropil. The lobula is associated with the lobula plate, which is wedged between the lobula and medulla on the posterior side of the optic lobe (Figure 3a, h). Both neuropils combined form the lobula complex, the final processing stage of visual information in the optic lobe. Like the lobula, the lobula plate possesses three layers, of which the posterior-most layer shows no 5HT immune-reactivity. Finally, the small accessory medulla is located near the medial rim of the medulla, directly adjacent to the anterior margin of the lobula (Figure 3a, c, d). This neuropil has no obvious internal structure and is approximately spherical in shape. Due to numerous protrusions towards the dorsal and ventral sides it has a rather irregular appearance.

**FIGURE 3.**
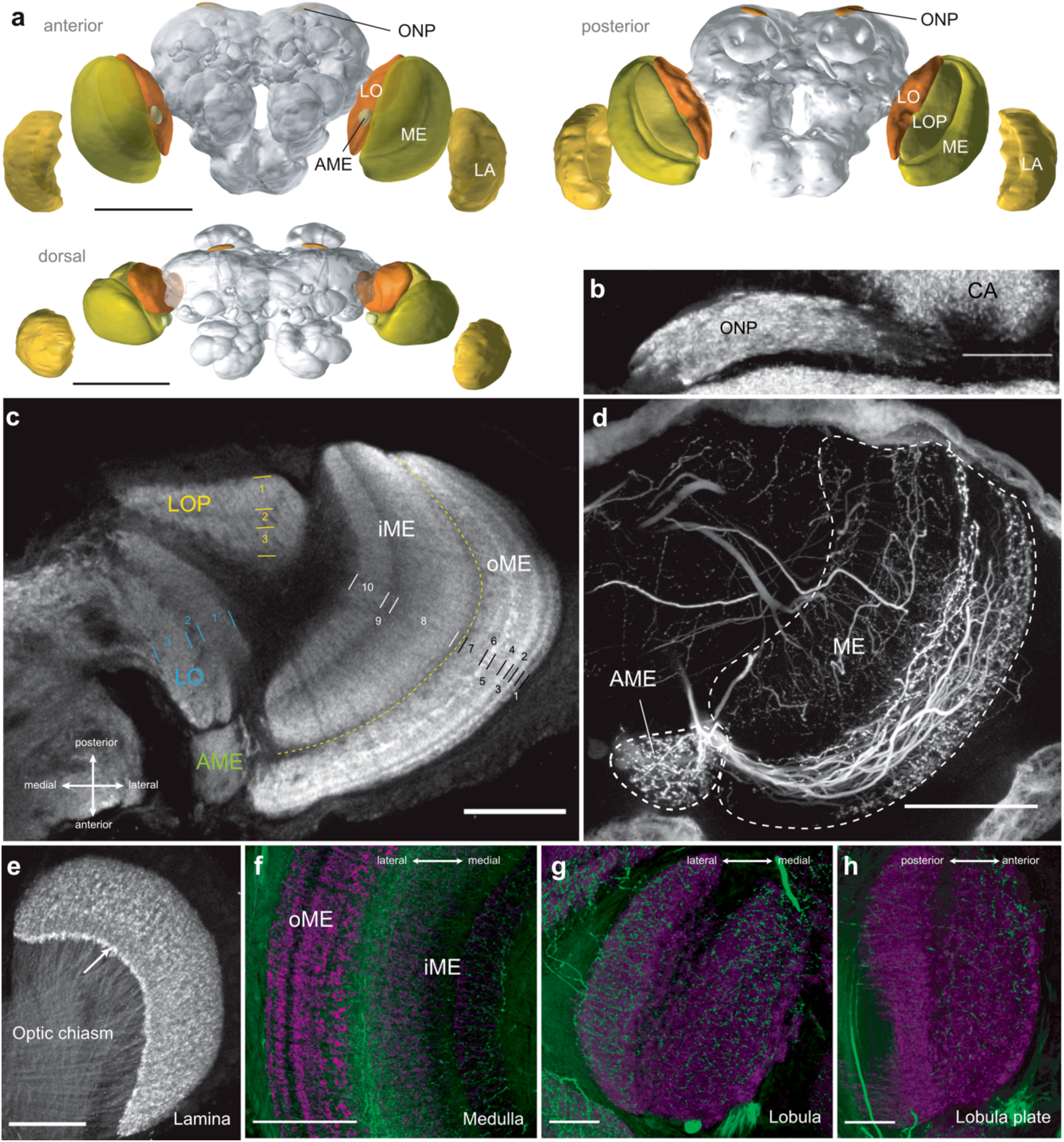
Primary visual neuropils. (a) Anterior, posterior and dorsal views of a 3D reconstruction showing the optic lobes including lamina (LA), medulla (ME), accessory medulla (AME), lobula (LO), lobula plate (LOP) and ocellar neuropils (ONP). (b) Left ocellar neuropil (ONP) stained against synapsin. These small neuropils are located near the dorsal surface of the Bogong moth brain and receive direct input from the ocelli via the ocellar nerve (not shown). (c) Horizontal section of the optic lobe, stained against synapsin. The 10 layers of the medulla (labelled white) are shown: the first seven layers comprise the outer medulla (oME) while layers eight to ten form the inner medulla (iME). iME and oME are separated by the serpentine layer. The lobula (LO; blue) and lobula plate (LOP; yellow) each show three distinct layers. The accessory medulla (AME; green) is located at the anterior end of the medulla. (d) Neurobiotin injection into the optic stalk shows a small subset of neurons that branch within the accessory medulla, while most other neurons branch in the medulla. (e) anti-synapsin staining of the lamina (LA), which clearly shows the inner, synapsin-rich layer. Neurons from the lamina connect to the medulla and cross over in a clearly visible optic chiasm. (F-H) Distribution of serotonergic fibres (stained with anti-5HT, green) within the medulla (e), lobula (f) and lobula plate (g). Neuropils are labelled with anti-synapsin (magenta). The serpentine layer (c) separates the inner and outer medulla, and most 5HT-positive neurons invade the medulla in this layer. Scale bar (a) 500 *µ*m, (b, f-h) 50 *µ*m, (c-d, e) 100 *µ*m.

The ocelli are connected to the brain via the thin ocellar nerve, which enters the brain dorsally near the mushroom body calyx. The axons of the ocellar photoreceptors terminate in the ocellar neuropil (ONP), an elongated, irregularly shaped brain area on either side of the midline (Figure 3a, b).

### Anterior optic tubercles

The target of most optic lobe projection neurons in insects are the optic glomeruli spread throughout the ventrolateral protocerebrum (Ito et al. 2014). The most prominent set of these glomeruli, as defined by Ito et al. (2014) in *Drosophila*, is a group of well-defined neuropils near the anterior surface of the brain. Combined, these neuropils are called the anterior optic tubercle (AOTU, Figure 4) and represent the first processing stage of visual information reaching the central brain from the optic lobes via the anterior optic tract. In the Bogong moth the AOTU is the only visible optic glomerulus and its overall layout was presented in de Vries et al. (2017). Consistent with this study, we found that the AOTU is located directly underneath the anterior surface of the Bogong moth brain, dorsally of the antennal lobes and laterally of the mushroom body lobes. It consists of three subunits: the upper unit, the lower unit and the nodular unit (Figure 4b-e). The nodular unit and the lower unit together form the lower unit complex. Whereas the lower unit had a homogeneous appearance in synapsin labelling, the nodular unit was further subdivided into four compartments. Dye injections into the optic lobe revealed that subsets of the four nodular-unit compartments are innervated by distinct optic-lobe projections. Similarly, the large upper unit receives projections that are not all homogeneously spread across the entire region, but innervate distinct ventral and dorsal aspects of this neuropil.

**FIGURE 4.**
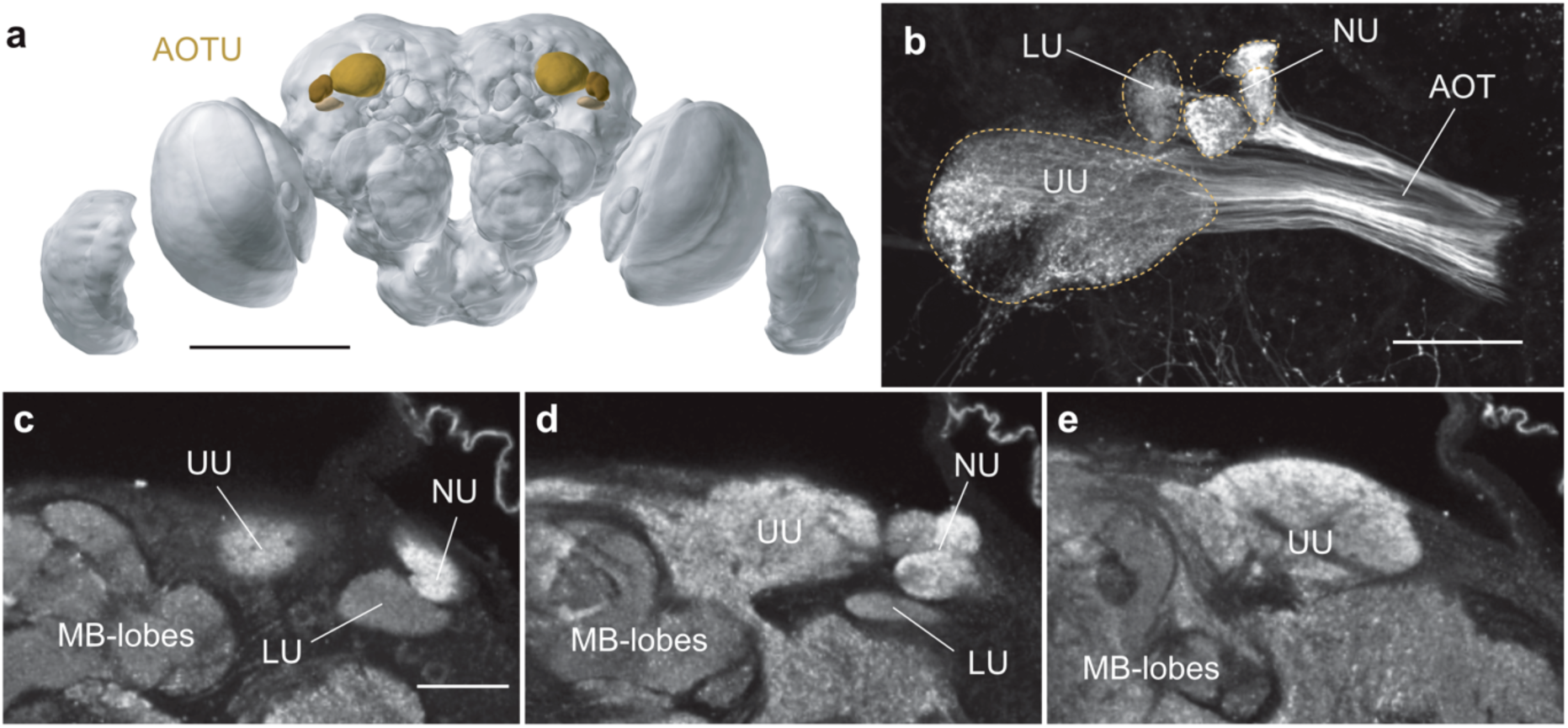
The anterior optic tubercle. (a) 3D reconstruction of the anterior optic tubercles (AOTU) in the Bogong moth brain. (b) Neurobiotin tracer injection into the optic lobe labels the upper unit (UU), lower unit (LU) and three subunits of the nodular unit (NU) of the AOTU. One subunit of the nodular unit is missing. AOT = anterior optic tract. (c-e) Optical horizontal sections through the AOTU show the upper unit, the full extent of the lower unit, and all four subunits of the nodular unit. Scale bar: (a) 500 *µ*m, (b, c) 50 *µ*m.

### Antennal lobes

The antennal lobes (AL, Figure 5) receive direct input from the antennae and are the primary processing stages for olfactory information. They are situated at the anterior margin of the central brain and are subdivided into individual processing units known as olfactory glomeruli. The glomeruli occupy the periphery of the AL and surround a central neuropil region that mostly contains fibres without anti-synapsin labelling. Each glomerulus is also characterized by a center-surround structure, highlighted by 5HT labelling that is restricted to the core of each glomerulus (data not shown). The periphery of the glomeruli shows bright anti-synapsin labelling. In total, the Bogong moth AL contains 72-75 glomeruli (six ALs from four moths). Near the anterior side of the AL, glomeruli are of regular, round appearance, whereas they become more irregular in shape in posterior regions of the AL, in particular towards the medial edge (Figure 5b). A group of well-defined, large glomeruli near the dorsal posterior side of the AL stood out as a distinct cluster.

**FIGURE 5.**
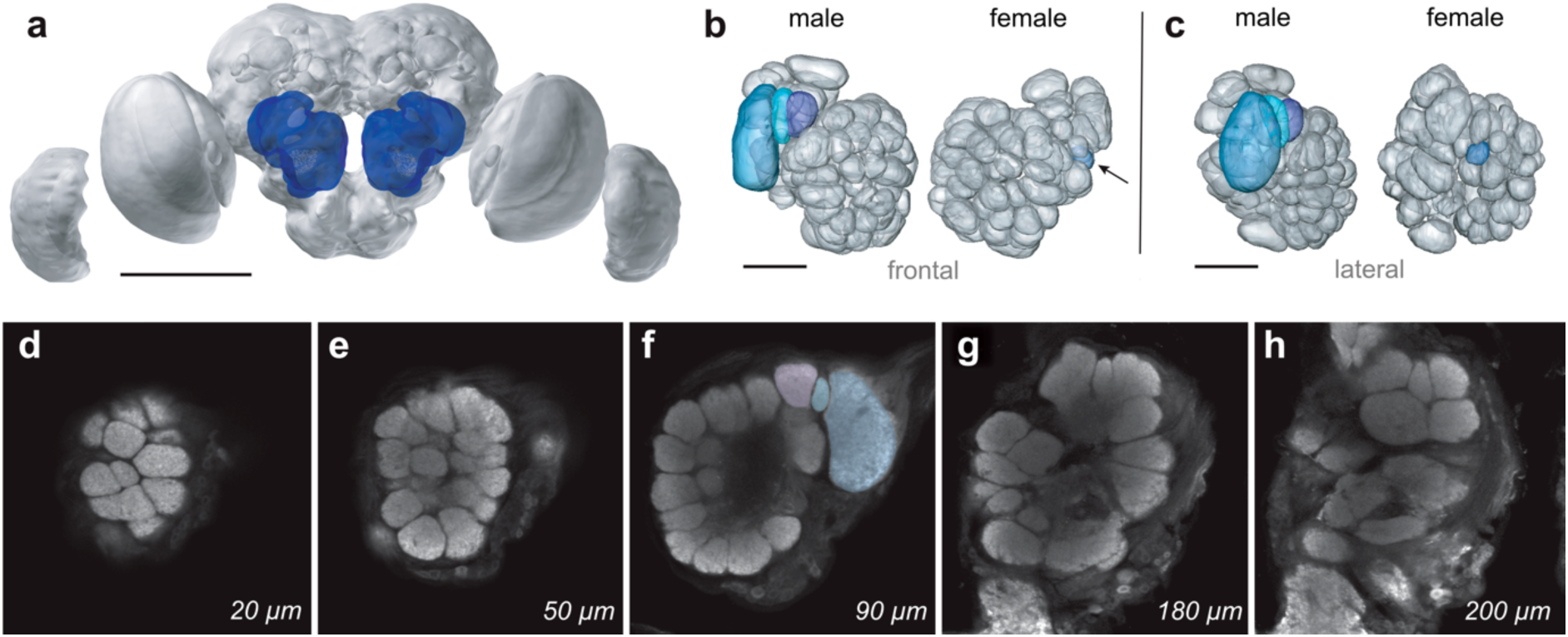
The antennal lobe. (a) 3D reconstruction of the Bogong moth brain with the antennal lobes (ALs) highlighted in blue. Scale bar = 500 *µ*m. (b) Detailed 3D reconstruction of a male and female AL. Out of a total of 72 glomeruli in the male AL, three glomeruli were identified as macroglomeruli (highlighted in blue, turquoise and purple) and constitute the macroglomerular complex (MGC). Female ALs have 73 glomeruli, including one that appears to be female-specific (highlighted in blue, arrow). (c) In both males and females, anterior glomeruli appear more regularly shaped than posterior glomeruli, which are also larger. Scale bar = 100 *µ*m. (d-h) Optical sections progressing from anterior to posterior through an anti-synapsin labelled male antennal lobe. The numbers indicate the depth level of each section, with 0 *µ*m marking the anterior border of the AL. The antennal nerve enters the AL near the cumulus of the MGC.

The ALs are the only neuropils of the Bogong moth brain that show an obvious sexual dimorphism. Three large male-specific glomeruli form the macroglomerular complex (MGC) near the entry point of the antennal nerve (Figure 5b, e). In females, the glomeruli in the same region are much smaller and cannot be distinguished from the ordinary non-MGC glomeruli. All well-defined, non MGC glomeruli could be individually matched between male and female preparations (n = 2 each). However, we found one glomerulus in females that had no identifiable counterpart in males and thus appears to be female-specific (Figure 5b). Whether this glomerulus is indeed female specific, or whether this region of the AL shows inter-individual variability with respect to glomeruli numbers remains to be shown.

### Mushroom bodies

Olfactory information leaves the AL via one of several tracts that transmit information to higher brain centers. The most prominent of these second-order olfactory regions is the calyx of the mushroom body, which together with the pedunculus and a complex system of lobes comprises a large, paired neuropil group, the mushroom bodies. Whereas the calyx is located at the posterior brain border, the lobes occupy the anterior part of the central brain, just dorsally of the ALs (Figure 6a). The pedunculus forms the connection between both parts along the antero-posterior brain axis, comprising dense bundles of axons that span the entire central brain. Input from the AL reaches the mushroom body in its posterior-most compartment, the calyx. One calyx is located on each side of the brain and each consists of an outer and an inner calyx, that is, two ring-like neuropils that are fused in the center, close to the origin of the pedunculus. Two holes are visible in the center of both rings in anti-synapsin labelled brains and are formed by bundles of primary neurites of Kenyon cells, the principal intrinsic neurons of the mushroom body (Sjöholm et al., 2005).

**FIGURE 6.**
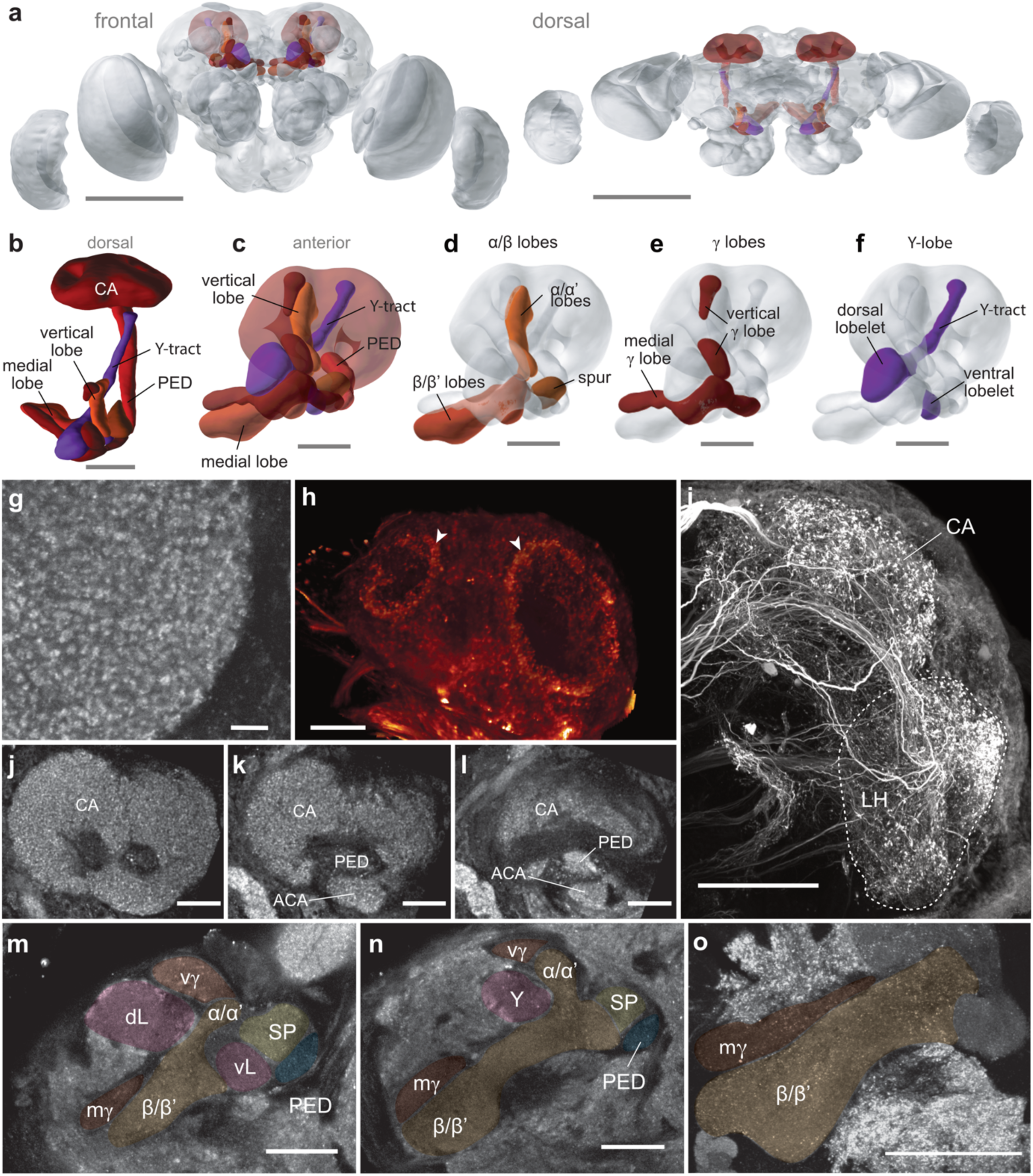
The anatomy of the mushroom body. (a) 3D reconstruction of the mushroom bodies (MBs) within the brain, frontal and dorsal view. The MBs span from the frontally-located lobes to the dorsal calyces. (b) Lateral view of the MB. (c) Frontal view of the MB. (d-f) Frontal view of the MB, with the three lobe systems highlighted. Scale bar = 100 *µ*m. (g) The calyx is composed of clearly visible micro-glomeruli. Scale bar = 10 *µ*m. (h) Injection of Neurobiotin into the optic stalk highlights neurons with output in the inner ring of the calyx (volume rendering of confocal image stack). (i) Neurobiotin injection into the antennal lobe labels projection neurons to the calyx of the MB and the lateral horn (LH). (j-l) Frontal optical sections (10 *µ*m steps) through the calyx reveal the peduncle and accessory calyx. (m) Frontal section through the MB lobe systems at a depth of 90 *µ*m. my = medial gamma lobe, ß/ß’ = medial beta lobes, vy = vertical gamma lobe, dL = dorsal lobelet of the y-lobe, vL = ventral lobelet of the y-lobe, Sp = spur, Ped = peduncle. (n) Frontal section through the MB lobe systems at a depth of 130 *µ*m. a/a’ = medial alpha lobes. (o) Frontal optical section through the medial beta/beta’ lobe and medial gamma lobe (maximum intensity projection over 10 steps, step size = 1 *µ*m). Scale bars in A = 500 *µ*m, B-F, G, M = 100 *µ*m, H-L, O-P = 50 *µ*m.

The calyces are characterized by small, globular synaptic domains that are evident from anti-synapsin labelling (Figure 6g). These microglomeruli are the terminal branches of input neurons to the calyx. Dye-injections into the AL revealed that they belong predominantly to olfactory projection neurons (Figure 6i). However, dye injections into the optic stalk demonstrated that a small set of projections originates not in the AL but in the optic lobe. Surrounding the two central holes in each hemi-calyx, these projections form a distinct inner ring that is nested within a larger outer ring, giving rise to an overall structure that consists of two sets of concentric neuropil rings in each calyx (Figure 6h). Tightly associated with the calyx is the accessory calyx, which also receives visual projections (Figure 6k, l). The accessory calyx is evident as a ventral protrusion from the calyx-proper, directly adjacent to the pedunculus. The borders of this region are not sharply defined and the neuropil is fused posteriorly with the calyx and anteriorly with the protocerebral mass that surrounds the pedunculus on its ventral side.

The pedunculus contains the axons of the Kenyon cells and links the input region of the mushroom body (calyx) to its principal output regions, the lobes. In the Bogong moth two primary lobe systems can be distinguished in synapsin-labelled preparations: the vertical and the medial lobes, each of which can be separated into two parallel sub-lobes. These two systems originate from the anterior end of the pedunculus and stretch medially towards the midline (medial lobe) and dorsally towards the dorsal brain border (vertical lobe; Figure 6c). The medial lobe contains the beta lobe and medial gamma lobe, while the vertical lobe contains the alpha lobe and the vertical gamma lobe (Figure 6d-e). Towards their distal ends, the beta lobe as well as the alpha lobe further separate into two parallel neuropil streams, likely corresponding to the alpha and alpha-prime, and the beta and beta-prime compartments known from other species (e.g. Sjöholm et al., 2005). Both lobe systems have a complex, convoluted structure towards the anterior brain surface, where the two arms of the gamma lobes meet and wrap around the roots of the beta and alpha lobes. In particular the ventro-lateral regions of the medial lobe have numerous protrusions and sub-lobes and it is not always obvious where the exact border between the alpha lobe and the overlaying gamma lobe is. Near the point where the vertical and medial lobes have their common root, we found another small neuropil that could not be assigned to one of the two lobes. This spherical region was called the spur and might indeed correspond to the spur region in *Drosophila* (Ito et al. 2014; Figure 6m, n).

Finally, as other lepidopteran insects, the Bogong moth also has a secondary lobe system with a separate connection to the calyx, the Y-tract (Figure 6f). This tract originates in dorsal regions of the calyx and runs parallel to the pedunculus towards the anterior side of the brain. It turns medially just posteriorly of the vertical lobe and supplies two lobelets that intermingle with the main lobe systems (Figure 6m, n). The ventral lobelet of the Y-lobe is comparably small and sharply defined. It protrudes through the beta lobe towards the ventral margin of the lobe system, where it borders the AL and the lateral accessory lobes (LAL). The dorsal lobelet is much bigger and fills large portions of the space stretching in between the vertical and medial lobes, expanding all the way to the anterior brain surface. This lobelet corresponds to the largest of three ellipsoid mushroom body lobe protrusions that are visible as landmarks on the brain surface. The remaining two are formed by the vertical and medial gamma lobes.

The second major target region for olfactory projections from the AL is the lateral horn (Figure 6i). The neurons terminating in the lateral horn either originate in the medial antennal lobe tract and reach the lateral horn after providing input to the calyx, or they target this region directly via the lateral and mediolateral antennal lobe tracts (Ian et al., 2016). The lateral horn neuropil is located in the lateral-most part of the central brain and is medially completely fused with the surrounding protocerebral regions (inferior, superior and ventrolateral protocerebrum). Dye injections into the AL revealed the location and approximate size of this area (Figure 6i), but further analysis will be required to delineate the exact borders of the lateral horn.

### Central complex

Together with the AOTU and the lateral complex, the central complex (CX) of the Bogong moth brain was examined in a previous study (de Vries et al., 2017) in the context of potential roles of these neuropil groups for migratory behavior. As this study was focused on identifying volumetric correlates of migratory lifestyle, it did not describe the detailed neuroanatomical features of the Bogong moth CX, which was therefore the aim of the current study.

The CX is a midline-spanning group of neuropils, located dorsally of the esophageal foramen. It consists of four compartments: the protocerebral bridge (PB), the fan-shaped body (FB; or central body upper division, CBU), the ellipsoid body (EB; or central body lower division, CBL) and the paired noduli. While the fan-shaped body and ellipsoid body are the only unpaired neuropils in the brain (Figure 7a, b), both the PB and the noduli occur in symmetrical pairs on either side of the midline. The EB is the anterior-most CX compartment and has a bar-like shape, with its ends bent slightly anteriorly. It spans the width of the central body from right to left and has clearly defined borders on all sides. The FB is directly adjacent to the EB on its posterior side and partially encompasses the EB from behind. It is approximately three times as large as the EB and has a pronounced stratified appearance (Figure 7d-g). Posterior to the FB, at the posterior border of the brain, lies the PB. The FB and the PB are separated by a stretch of neuropil belonging to the inferior protocerebrum that possesses two large openings on either side of the midline, allowing the passage of neurites be-tween the two CX compartments. Each half of the PB is visible as a brightly synapsin-labelled, hook-like structure (Figure 7k). Both emerge near the midline, bend dorsally, and then continue ventro-laterally in a straight line, approximately to the level of the mushroom body pedunculus. The medial parts of the PB have a larger diameter, which continually decreases towards the lateral ends. The final lateral segment of the PB often stands out as a distinct, globular domain. A thick fiber bundle interconnects the right and left PB across the midline. The final CX neuropils, the noduli (NO), are located ventrally of the FB and the EB. These neuropils are the smallest part of the CX and consist of a large and a small compartment, both of which are separable into two layers (Figure 7h-j).

**FIGURE 7.**
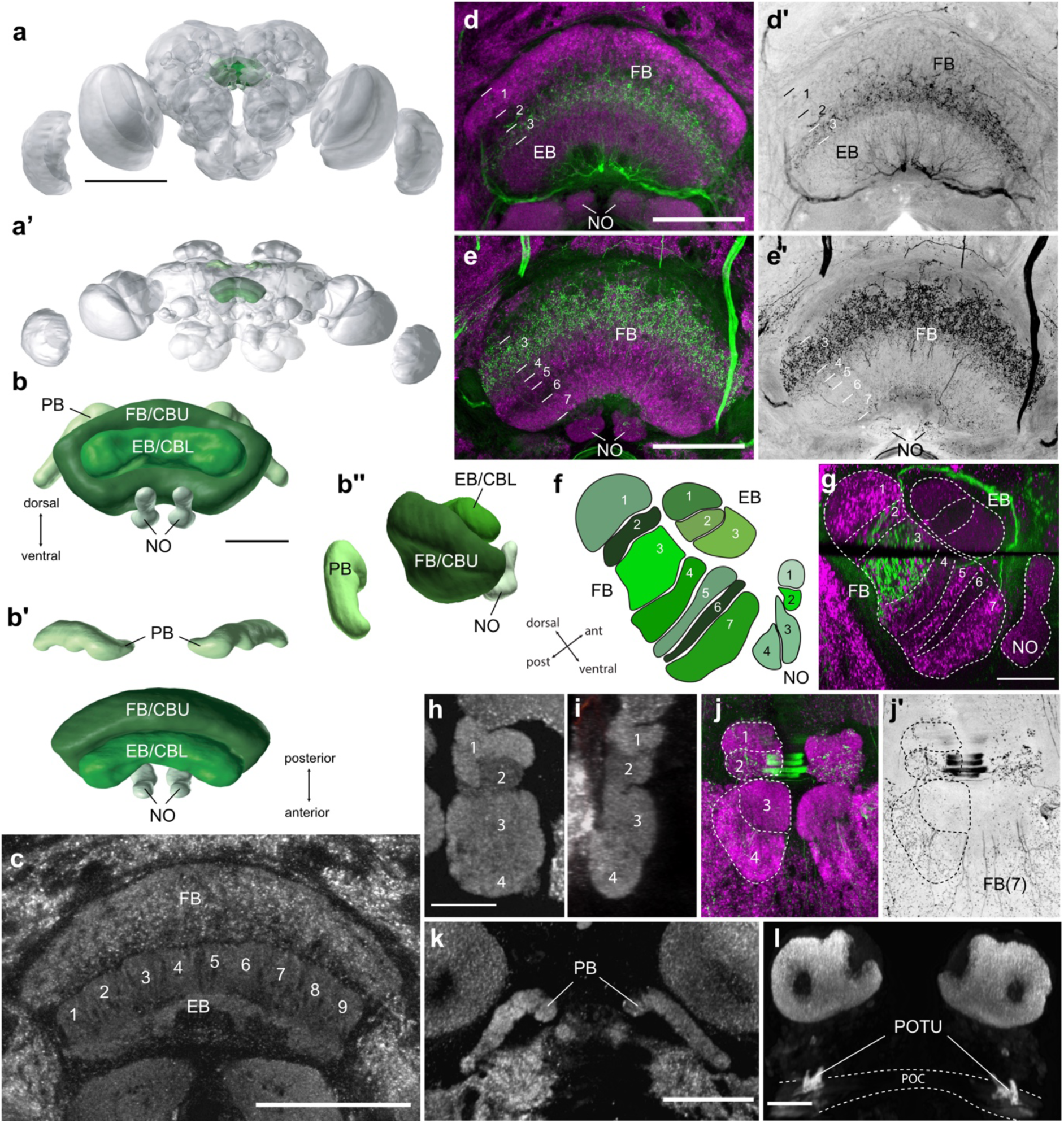
The morphology of the central complex. (a) Frontal view of a 3D reconstruction of the Bogong moth brain, with the central complex (CX) highlighted in green. Scale bar = 500 *µ*m. (A’) Dorsal view of the same reconstruction. (b-b’’) 3D reconstruction of the CX, frontal view (b), dorsal view (b’), and lateral view (b’’). Scale bar = 100 *µ*m. (c) Frontal confocal section of the CX, highlighting the ellipsoid body (EB and its clearly visible lateral slices. Scale = 100 *µ*m. (d) Anti-synapsin (magenta) and anti-5HT (green) staining highlight the layers of the fanshaped body (FB). Shown is a maximal intensity projection (depth 5 *µ*m) at the level of the EB and layer 1-3 of the FB. Scale = 100 *µ*m. (d’) 5HT labelling from D, isolated and shown with inverted colormap. (e, e’) As D and D’, but for a cross-section at a more posterior level, showing layers 3-7 of the FB as well as 5HT labing in the noduli (NO). (f) Schematic cross-section of the central body and noduli. The FB has seven layers defined via anti-synapsin and anti 5HT labelling. The three layers of the EB are visible as well as all four subunits of the noduli. (g) Sagittal cross-section of FB showing the 7 individual layers (maximal intensity projection, 12 images, step size = 0.4 *µ*m); compound image from two physical thick-sections. The black gap across the FB is the border of the two slices. Scale = 100 *µ*m. (h) The paired noduli have four subunits each, visible in this horizontal confocal section. Scale bar = 50 *µ*m. (i) Sagittal view of the same preparation as in (h). (j) Maximal intensity projection of a preparation labelled against synapsin and 5HT, revealing the 5HT labelling in layer 2 of each nodulus. (J’) Isolated 5HT labelling from (j), highlighting the serotonergic innervation of layer 2, as well as the small, beaded terminals of FB layer 7 (partly superimposing the ventral layers of the noduli). (k) Frontal view of the protocerebral bridge (PB; maximal intensity projection, 10 images, step size = 1 *µ*m). Scale = 100 *µ*m. (l) Frontal view of the posterior optic tubercle (POTU). Maximal intensity projection (16 images, step size = 2 *µ*m) showing the POTU located posterior to the posterior optic commissure (POC). Scale = 50 *µ*m.

Given the complexity of the neuroarchitecture of the CX, the tight structure-function relations in this region across insects, and its likely role as an internal compass also in the Bogong moth, we will highlight the distinguishing features of each of the CX compartments. Most prominently, the FB exhibits a pronounced stratification that was revealed via a combination of synapsin labelling and anti 5HT labelling. Using these techniques as defining markers, this neuropil is subdivided into seven layers (Figure 7d-g). These layers are linearly stacked on top of each other from dorsal to ventral. Layer 1 is easily distinguished by its arch like morphology and by displaying the brightest synapsin labelling within the FB. Adjacent ventrally is layer 2 and 3, showing weaker synapsin labelling. Layer 3 is a thick, strongly serotonergic stratum, characterized by large, beaded 5HT positive terminals (Figure 7e). The overlaying thin layer 2 shows only weak 5HT labelling, which originates from 5HT-labelled fibres that pass through this layer. The ventral half of the FB contains weaker 5HT labelling visible in two out of the four remaining layers. We distinguished four layers (4, 5, 6 and 7) according to differences in the visual appearance of synapsin labelling and the presence of 5HT labelling (Figure 7e’, f). Unlike the FB, the second part of the central body, the EB, has a less pronounced stratification (Figure 7c,d). 5HT labelling was found evenly distributed in the EB but was less distinct than in the FB. Based on synapsin labelling, three layers were identified. The dorsal-most layer was distinguished from the remaining EB due to the presence of nine clearly delineated lateral divisions, i.e. slices (Figure 7c). A columnar structure was partially visible in the PB as well, but was more obvious in lateral parts of the neuropil. Towards the midline, columns fuse into a homogeneous neuropil without any internal features. Finally, the noduli consist of four layers each, two in the small dorsal compartment and two in the larger ventral compartment. The first layer forms a cap-like region of bright synapsin labelling, while the remaining three layers are less clearly delineated (Figure 7h-j). Layer 2 of the noduli shows sparse innervation by serotonin-immunoreactive beaded fibers (Figure 7j), which were labelled in varying intensity between individual preparations.

A region closely associated with the PB is the posterior optic tubercle (POTU; Figure 7l). While not strictly part of the CX, this irregularly shaped neuropil is located at the posterior surface of the brain on either side of the midline. It is connected to the PB via two thin fiber bundles, which extend beyond the length of the PB towards ventrolateral directions. The POTU is located at the intersection of this bundle and the horizontally running posterior optic commissure (POC, Figure 7l), a major commissure interconnecting the optic lobes on either side of the brain.

### Lateral complex

The final set of well-defined neuropils in the Bogong moth brain is the lateral complex. This brain area consists of three distinct compartments and is located immediately adjacent to the CX, flanking it laterally and ventrally on both sides. It stretches from the level of the central body towards the anterior brain surface and borders anteriorly with the AL and the mushroom body lobes. Each lateral complex consists of the large lateral accessory lobe (LAL), complemented by the smaller bulb and the gall, which are situated on the anterio-dorsal surface of the LAL (Figure 8a, b).

**FIGURE 8.**
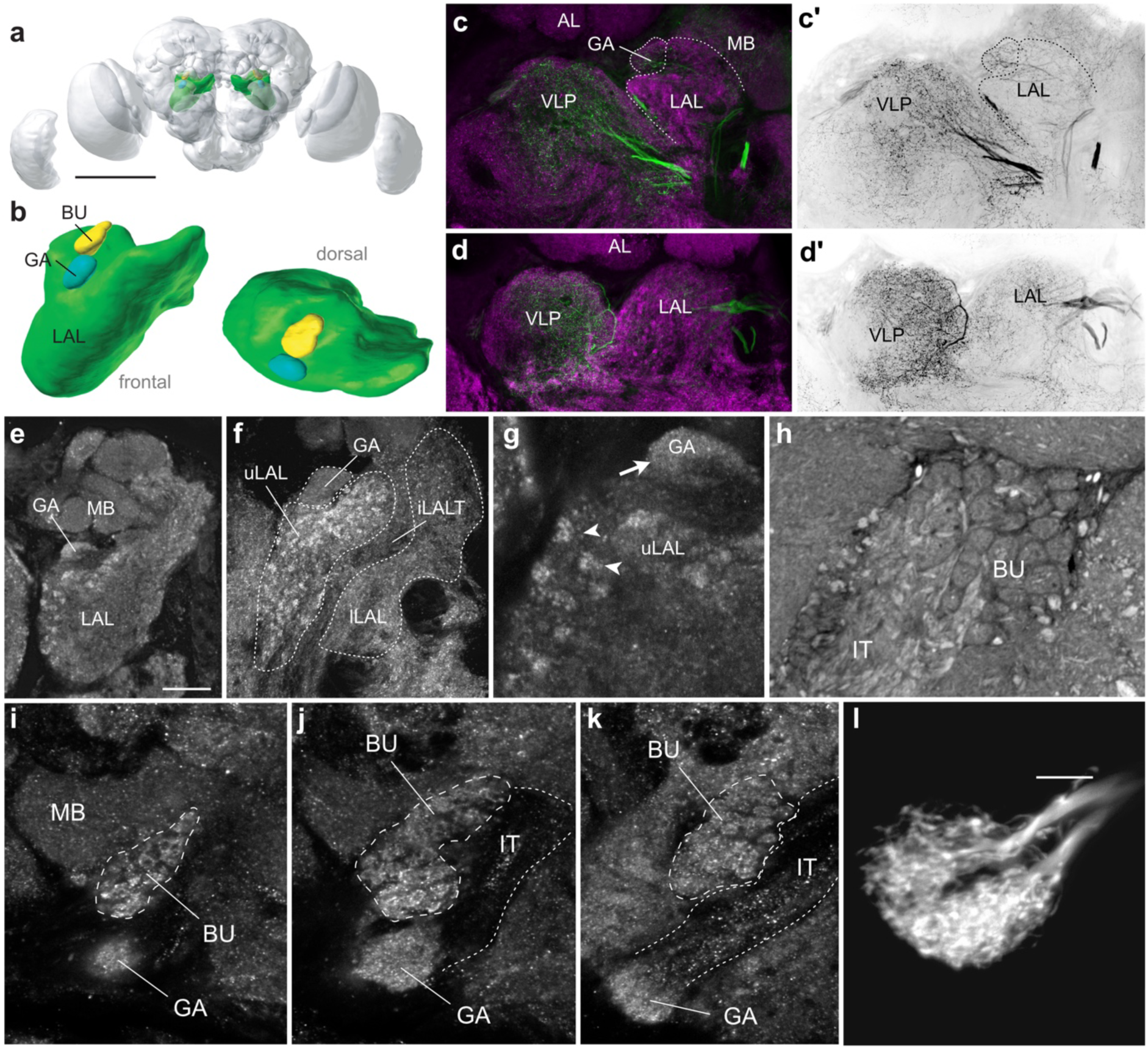
The morphology of the lateral complex. (a) 3D reconstruction of the lateral complex in the Bogong moth brain. Scale bar = 500 *μ*m. (b) Frontal and dorsal view of the lateral complex, with the lateral accessory lobes (LALs) in green, the bulb (BU) in yellow and the gall (GA) in turquoise. (c, c’, d, d’) Horizontal section of the LAL showing a clear boundary to the heavily serotonin-labelled ventrolateral protocerebrum (VLP). The two sections are 120 *μ*m apart. Anti-synapsin labelling is shown in magenta, anti-5HT staining in green. (e) The LAL lies immediately adjacent to the mushroom body lobes, with the gall nestled between the two neuropils. Scale = 50 *μ*m. (f) The inter-LAL tract (iLALT) subdivides the LAL into the upper LAL (uLAL) and the lower LAL (lLAL). Scale bar = 50 *μ*m. (g) Anti-synapsin stained microglomeruli in the dorsal shell of the LAL. Scale = 50 *μ*m. (h) Light microscope image of the bulb stained with Azur II-Blue, with individual microglomeruli visible. (i-k) Horizontal optical sections showing the bulb and gall lying on top of the dorsal shell of the LAL, sections progress from dorsal to ventral. Both neuropils are served by fibres in the isthmus tract (IT). Scale bar = 20 *μ*m. (l) Neurobiotin-labelled CL1 cell branching in the gall. The two subdivisions of the gall are clearly visible. Scale bar = 10 *μ*m.

The borders of the LAL are well defined along its anterior and medial margins. In those regions it is bordered by the AL (anteriorly) and the medial antennal lobe tract (medially). The lateral boundary is evident with the help of anti-5HT labelling, which highlights a pronounced difference in staining pattern and intensity between the bordering ventrolateral protocerebrum and the lateral portions of the LAL (Figure 8c, d). Dorsally, the bulb, the isthmus tracts and the mushroom body lobes create natural boundaries in most regions, which can be generalized into a dorsal cut-off plane for more ambiguous areas near the anterior brain surface and the posterior extremes of the LAL. The posterior boundary is defined by the horizontal crossing of the lateral antennal lobe tract behind the LAL, which defines the plane that separates the LAL from the more posterior regions of the ventromedial protocerebrum. While this border is somewhat arbitrary, it is consistent with the LAL definitions in other species (Heinze and Reppert, 2012; Ito et al., 2014; Immonen et al., 2017). More clearly defined than the LAL, the bulb is wedged between the dorsal LAL margin and the posterio-medial surface of the mushroom body lobes (Figure 8i-k). It provides a cap-like endpoint to the isthmus tracts, which are fiber bundles that emerge from the lateral edges of the EB. As in other insects, the bulbs are thus tightly linked to the EB. Just antero-ventrally of the bulb is the gall, which is served by a subset of isthmus tract fibers that continues beyond the bulbs.

The largest compartment of the lateral complex, the LAL is further subdivided into the upper LAL and lower LAL by the inter-LAL tract (Figure 8f), which connects the two lateral complexes. The two compartments differ in their internal structure, innervation patterns of serotonergic fibers, and extent of fusion with neighboring regions (Figure 8c, d). While the upper LAL contains numerous, irregularly shaped glomerular domains that light up brightly with anti-synapsin labelling (Figure 8g), no such internal structure was observed in the lower LAL. The borders of the upper LAL are more distinct than those of the lower LAL, which merges with the surrounding ventrolateral neuropils. Future functional data will thus have to show whether the borders of the lower LAL, although consistent with other insects, indeed represent a functional unit.

While the bulb has no easily discernible subunits, the gall contains two equally sized compartments (Figure 8i-l), each of which is targeted by a set of columnar neurons from the EB (CL1 or E-PG neurons; Figure 8l). These gall compartments have an appearance not unlike AL glomeruli, with a dark, synapsin-deficient core and a synapsin-rich rind. No finer scale structure is visible within these regions. This is in stark contrast to the bulbs, which exclusively consist of sharply defined, ball-like synaptic domains. These microglomeruli are highly visible in Azur II–Methylene Blue staining (Figure 8h), but are also apparent with synapsin immunolabeling, which is concentrated in ring-like domains surrounding the periphery of each microglomerulus (Figure 8i-k).

## Discussion

Bogong moths are remarkable long-distance navigators that are guided by a comparably simple brain. Understanding the neural circuitry of this brain will provide a unique window into the neural basis of nocturnal long-distance migration. With the current study, we have provided a morphological framework for future functional work that aims at understanding the neural basis of migration and navigation in the Bogong moth.

In general, the Bogong moth brain is similar to the brains of other lepidopteran insects, such as the Monarch butterfly *Danaus plexippus* (Heinze and Reppert, 2012), *Heliothis virescens* (Kvello et al., 2009), *Manduca sexta* (el Jundi et al., 2009), or the hummingbird hawkmoth *Macroglossum stellatarum* (Stöckl et al. 2016). Given the spectacular migratory lifestyle of the Bogong moth, it is noteworthy that there is no single gross morphological feature of the brain that stands out as unique or different compared to related species. In contrast, it shares not only all major features of the overall insect brain ground pattern, it also exhibits all distinguishing characteristics of other lepidopteran brains. These include a subesophageal ganglion that is fused with the brain, well-developed optic lobes, large antennal lobes, a complex organization of the mushroom body with a multi-layered double calyx and an intricately organized lobe system. One feature of the mushroom body in particular appears to be a defining characteristic of moths and butterflies, namely the presence of the Y-tract with its associated lobelets (Heinze and Reppert 2012; Sjöholm et al. 2005; Montgomery et al. 2016; Homberg et al. 1988). A second unusual feature of lepidopteran brains is that the PB of the CX is split across the midline. In other insects this neuropil is visible as a continuous, midline-spanning structure (Heinze and Reppert 2012; Stöckl et al. 2016; el Jundi et al. 2009).

With 25 separately segmented neuropils, our average atlas of the Bogong moth brain is the most detailed reconstruction of a moth brain to date (el Jundi et al. 2009; Kvello et al. 2009; Stöckl et al. 2016). It was carried out in exactly the same way as the reconstruction of the Monarch butterfly brain (Heinze and Reppert, 2012), thus offering the basis for detailed comparisons between a day-active migratory butterfly and a night-active migratory moth. In the following sections we will therefore aim at identifying morphological features that stand out as commonly shared between those two migrants, when compared to non-migratory lepidopterans - features that could indicate specializations due to a migratory lifestyle.

### Primary sensory brain areas

Two sensory modalities likely guide the long range flights of both the Bogong moth and the Monarch butterfly: visual cues and geomagnetic cues (Dreyer et al. 2018; Guerra et al. 2014). Additionally, olfactory information has been proposed to be decisive during the last segment of the migration, i.e. when locating the overwintering grounds or the aestivating caves (Warrant et al. 2016; Mouritsen et al. 2013). While the sensors of magnetic field cues are not known in any species, the antennae have been implicated in magnetosensation in the Monarch butterfly (Guerra et al. 2014). However, a vision-based mechanism to detect magnetic fields remains a main hypothesis across many animal species (Ritz et al. 2000; Schulten et al. 1978), including in the Bogong moth. In any case, the processing of visual information, geomagnetic information, and olfactory cues would involve both the antennal lobes and the primary visual brain centers.

In the Bogong moth antennal lobes we found a pronounced sexual dimorphism, highly reminiscent of other moths, but unlike the Monarch butterfly, in which both sexes showed an identical AL layout (Heinze and Reppert, 2012). In other moths, including the closely related *Agrotis segetum* and *A. ipsilon*, the male-specific part of the AL (the MGC) processes female pheromones (Hansson et al. 1994; Jarriault et al. 2010; Berg et al. 2002). Although the pheromone of the Bogong moth has yet to be identified, it is likely that pheromone-based communication exists in the Bogong moth and that the MGC serves the same function as in the other species. In the context of migration, Bogong moths might pinpoint their aestivation caves based on odor cues that were left by previous generations of moths, such as excrement on the cave walls, and moth debris on the cave floor (Warrant et al. 2016; Dreyer et al., in preparation). As both male and female moths are faced with this identical challenge, the MGC, as well as any female specific glomeruli, are not expected to play a role in this behavior. As the number of identified glomeruli corresponds well to other noctuid moths, e.g. *Helicoverpa armigera* (Zhao et al., 2016), we suggest that a direct, quantitative comparison of glomerular maps might be possible across species. The presented antennal lobe map thus provides a baseline for a detailed comparison of glomeruli not only between migratory and non-migratory life stages of Bogong moths, but also between this species and non-migratory moths. These analyses will help to locate potential neural specializations that mediate the ability of migratory lepidopteran insects to identify the highly specific, yet unfamiliar target sites of their migratory journey.

The primary visual areas of the Bogong moth brain are the optic lobes and the ocellar neuropils. Morphologically, the optic lobes of the Bogong moth have a structure similar to those of the Monarch butterfly (Heinze and Reppert, 2012) and indeed most other insects (Strausfeld 2012). As expected from the nocturnal lifestyle of the Bogong moth, the total relative volume of the optic lobe is about 50% smaller than in the diurnal Monarch butterfly, consistent with data on other butterflies (Montgomery et al. 2016). Interestingly, despite the overall size difference, the internal layout of each optic lobe neuropil is remarkably conserved between the two species. Using synapsin labelling as a marker, we found the same number of layers in all neuropils and the visual appearance of neuropils was highly consistent between the two species (Heinze and Reppert, 2012). Furthermore, serotonin labeling was also used to characterize the optic lobe neuropils in *Manduca sexta* (Homberg and Hildebrand, 1989). Whereas some differences in the details of the medulla layering are evident and the division of the lobula in an outer and inner lobula is more pronounced in *Manduca*, we found that the overall patterning of the neuropils is remarkably conserved. Note that the number of layers in the medulla (*Manduca*: 7; Bogong moth: 10) is not directly comparable, as one was defined based on serotonin labeling and the other based on synapsin labeling.

The second primary visual centers in the Bogong moth brains are the ocellar neuropils. These regions process input from the ocelli, the two small lens eyes located dorsolaterally on the moth’s head. While the Monarch butterfly also possesses very small lateral ocelli, as well as an associated neuropil near the dorsal brain surface (personal observations), no details about these structures have been described. Across insects, ocelli have diverse functions: In arctiid and noctuid moths, they mediate diurnal activity patterns by measuring ambient light levels (Eaton et al. 1983; Wunderer and de Kramer 1989), in dragonflies they play a role in flight control (Stange, 1981), and the ocelli of several nocturnal bees and wasps can detect polarized light, suggesting a possible role in navigation (Zeil et al. 2014). A potential role of lepidopteran ocelli for detecting migration-relevant cues cannot be excluded, but given their small size, visual information would likely be limited to sensing illumination levels and, potentially, spectral composition and polarization angles of skylight cues.

An alternative role for both the optic lobe and the ocellar neuropils arises from the finding that the Bogong moth is able to perceive the geomagnetic field (Dreyer et al., 2018). The location of the magnetic sense and the identity of the sensor are unknown, but similar to proposals in migratory birds, magnetic information might be detected in the retina via a radical-pair based mechanism mediated by a blue-light receptor molecule called cryptochrome (Schulten et al. 1978; Ritz et al. 2000). Beyond the retina, the optic lobes are thus convenient access points to study neural responses to changes in the magnetic field, as are the ocellar retinae and their associated neuropils.

Overall, while all primary processing stages for visual and olfactory information are highly developed in the Bogong moth, we found no obvious specializations that would suggest a specific role during migration. Similarities and differences between species appear to be linked to the diurnal versus nocturnal lifestyle and the location of these species on the lepidopteran phylogenetic tree. While a potential role in magnetoreception for the visual neuropils is a fascinating idea, we could neither support nor dismiss this possibility based on our anatomical data.

### Anterior optic tubercles

In the Bogong moth, the AOTU is the only anatomically distinct direct target region of optic lobe projection neurons. While this region is present in all insects that have been studied to date, its morphological composition is less conserved compared to the optic lobes. Generally, the AOTU consists of two main subunits: a large upper unit and a smaller, more diverse lower unit complex (LUC). The LUC can consist of only one region (e.g. in locusts: (Homberg, Hofer, Pfeiffer, & Gebhardt, 2003), or consist of several components with more or less defined internal borders (in bees: Mota et al. 2011; Zeller et al. 2015; in dung beetles: Immonen et al. 2017).

Within the Lepidoptera, a large upper unit and two main LUC compartments (lower unit and nodular unit) are present, but species differ in the specific composition of these subunits (el Jundi et al., 2009; Heinze and Reppert, 2012). While the nodular unit of the Bogong moth AOTU has four compartments, only three were found in the Monarch butterfly (Heinze and Reppert, 2012). Furthermore, the lower unit of the Monarch butterfly extends into an elongated structure called the strap, which is not present in the Bogong moth. As the AOTU of the butterfly *Godyris zavaleta* possesses an internal composition similar to that of the Monarch butterfly (Montgomery et al., 2016), the differences in LUC structure are either due to phylogenetic constraints or driven by higher visual demands in response to the butterfly’s day-active lifestyle. The finding that the AOTU of the diurnal hawkmoth *Macroglossum stellatarum* was very similar to that of other moths (Stöckl et al. 2016) points to phylogeny as the most important factor determining the differences within lepidopteran insects. Detailed investigations comparing size differences in the various AOTU divisions revealed a consistently larger upper unit in day-active species (Stöckl et al. 2016; de Vries et al. 2017). Additionally, a smaller lower unit was found in the day-active migrant *M. stellatarum* compared to the nocturnal non-migrant *Deilephila elpenor* (Stöckl et al. 2016), while a larger nodular unit was found in the Bogong moth compared to its close relative *Agrotis segetum* (de Vries et al., 2017).

Functional studies in the desert locust and Monarch butterfly showed that the AOTU-LUC is important for processing information about polarized light and the position of the Sun (Pfeiffer et al. 2005; Pfeiffer and Homberg 2007; Heinze and Reppert 2011). For these long distance migratory insects, polarized skylight serves as a compass cue used to determine the insects’ heading. Their AOTU contains at least two parallel pathways passing through the different regions of the LUC, carrying physiologically distinct information (Heinze, 2014). In *Drosophila*, two similar parallel pathways encode different aspects of landmark information (Omoto et al., 2017; Shiozaki and Kazama, 2017). In all species, these pathways share the bulbs of the lateral complex as their sole target and thus provide input to the head-direction system of the CX, suggesting that this is also the case in the Bogong moth.

Which sensory information can be used as a direction signal for the nocturnal Bogong moth? Polarized light from the sun is available until the sun is 18° below the horizon, thus until approximately one hour after sunset (Cronin et al. 2006). Aestivating Bogong moths have been observed to swarm in the hour after sunset, and they appear to depart on their migration within this time as well (Common, 1954). Thus, polarized light from the sun may be a viable directional cue during the first hour of flight. Later at night, polarized light from the Moon can also be used for course control, and possibly the Moon’s position itself as well as the Milky Way (Warrant and Dacke 2016). Independent of which compass cues are used for migration in the Bogong moth, all are likely relayed via the AOTU, as this is the only currently known pathway providing allothetic sensory input to the head direction system of the CX. The AOTU is therefore a primary target for physiological studies of the sensory basis for migratory behavior in the Bogong moth.

### Central and lateral complex

The CX is the brain region most heavily involved in spatial orientation and navigation, and it is therefore likely crucial during the Bogong moth’s migration (de Vries et al., 2017). It is anatomically conserved throughout the insects, likely because of its fundamental function in spatial orientation. As in all insects, the Bogong moth CX is characterized by horizontal layers and vertical slices, which are most visible in the EB. This arrangement is identically to that in the locust (Williams, 1975) and the dung beetle (el Jundi et al., 2019). Across many insects, the columnar structure results from a highly conserved underlying neural organization (Heinze and Homberg, 2008; Heinze et al., 2013; Wolff et al., 2015; el Jundi et al., 2019). Columnar neurons form the basis of a neural circuit that that encodes heading direction (Green et al., 2017; Turner-Evans et al., 2017). In two other long-distance migrating insects, the Monarch butterfly or the desert locust, this circuit uses the Sun’s position and the skylight polarization pattern to encode heading in a global reference frame (Heinze and Reppert 2011; Homberg 2015). In flies and cockroaches, local visual landmarks are used for this task (Seelig and Jayaraman, 2013; Varga and Ritzmann, 2016), but idiothetic cues can additionally provide information to compute a heading signal in darkness (Turner-Evans et al., 2017). The high degree of functional and anatomical conservation of this circuit across a wide range of insects suggests that the corresponding cells in the Bogong moth CX also encode heading in this species.

Beyond knowing the heading, the moths also have to know their target direction during the migratory journey. Based on a computational model that describes the CX as a neural substrate for path integration (Stone et al., 2017), it was recently suggested that this network could be modified to produce migratory behavior (Honkanen et al., 2019). The proposed model contains a set of memory neurons whose combined activity encodes the home vector in a path-integrating insect, generating an activity peak that signals the current nest direction and which is used by a steering cell population to keep the insect on track. While this activity peak dynamically changes with the ever-changing home vector, a migratory heading needs to be fixed and genetically encoded. This could in theory be accomplished by fixing the synaptic weights of the neurons corresponding to the memory cells, so that their population generates a constant output signal causing the animal to steer into the migratory heading whenever it is moving (Honkanen et al., 2019). Although experimentally challenging, the predicted activity patterns as well as the potential synaptic weight distributions are testable, and the detailed descriptions of the brain regions involved that we have provided in our current study lay the groundwork for these studies.

Finally, the CX gives direct output to the lateral accessory lobes (LALs), a brain region that has been implicated in initiating behaviors. Pre-motor neurons descend from this region into the ventral nerve cord and directly control steering in the silkworm moth (Namiki and Kanzaki, 2016). The Bogong moth LAL is anatomically very similar to that of the silkworm moth (Iwano et al., 2010). Furthermore, neurons that were described to play a role in phonotactic steering in crickets (Zorovic and Hedwig, 2011) closely resemble the ones involved in steering in the silkworm moth, suggesting that the circuit is indeed widely conserved. The Bogong moth LALs are therefore a likely substrate for a similar steering circuit, which can now be probed by electrophysiological recordings to illuminate how the moths are able to steer precisely towards their migratory target over many hundreds of kilometers.

### Mushroom bodies

Besides the CX, the mushroom bodies (MBs) are the second major integration center in the insect brain. This region is tightly associated with olfactory learning and memory (Strausfeld et al. 1998), but in many species also receives input from other modalities, such as vision (Gronenberg 2001; Lin and Strausfeld 2012; Paulk and Gronenberg 2008). In principle, the mushroom body associates stimuli with a reward signal and thereby gives sensory information a valence based on previous experience (Strube-Bloss et al., 2011; Aso et al., 2014; Menzel, 2014). The Bogong moth mushroom bodies are similar to those described in other lepidoptera (Strausfeld et al., 1998; Sjöholm et al., 2005; Sinakevitch et al., 2008; Heinze and Reppert, 2012; Montgomery and Ott, 2015), suggesting similar roles in learning and memory.

In other lepidopteran insects the input to the mushroom body is multisensory and the calyx consists of an olfactory outer and a non-olfactory, often visual, inner region (Stöckl et al. 2016; Sjöholm et al. 2005; Homberg et al. 1988; Kinoshita et al. 2015). In the Bogong moth, the inner ring of the calyx and the accessory calyx are indeed supplied with visual input, while the large outer ring is supplied by olfactory inputs. Whereas the sensory projections are not clear for the Monarch butterfly, the outer and inner zones of the calyx also exist in this species, but are complemented by the basal zone, giving the calyx of the Monarch butterfly a more complex appearance (Heinze and Reppert, 2012). Differences between the Bogong moth and the Monarch butterfly are also apparent in the lobe system, which in the Bogong moth closely resembles that of other moths, but has an unusual morphology in the Monarch butterfly (Heinze et al. 2012). Overall, complexity of lifestyle and sensory preferences of species have been shown to affect the structure of the mushroom body across insects (Molina and O’Donnell, 2008; Amador-Vargas et al., 2015). As migration in an inherited direction, while challenging, is neither complex nor flexible, the mushroom bodies are likely not related to compass-based navigation, but instead crucial for learning in the context of odor- and color-based foraging. In line with this idea, the observed differences in this brain region between the Monarch butterfly and the Bogong moth suggest that migratory behavior as such puts no constraint on the mushroom body.

Nevertheless, as outlined above, odors are likely of key importance for Bogong moths to locate their aestivation caves and could provide the signal that terminates the migration (Warrant et al. 2016; Dreyer et al., in preparation). In fruit flies, neurons of the lateral horn encode odor valence (Strutz et al., 2014; Schultzhaus et al., 2017) and specifically underlie innate attraction or aversion to behaviorally highly relevant odors (Jefferis et al., 2007). The Bogong moth lateral horn is therefore an interesting target region for functional studies that aim at understanding the mechanisms underlying the final segment of migration – the short distance search for their aestivation caves.

## Conclusion

The Bogong moth is a fascinating example of a nocturnal long-distance migrating insect. The anatomical framework of the brain presented here provides a basis for qualitative and volumetric comparison between the brains of Bogong moths in different migratory states, as well as for interspecific comparisons to migratory and non-migratory insects. Second, we can now target specific brain regions with electrophysiology or molecular methods in order to explore their role in encoding compass information and guiding migratory behavior. Finally, anatomical data from identified neurons, obtained from dye injections in conjunction with electrophysiological recordings, can be analyzed more extensively by registering the neurons’ anatomies into our standard atlas and thereby delineating functional neural circuits. Thus, by enabling future functional studies to be embedded within a detailed anatomical framework, we facilitate insights into the neural networks that guide all aspects of migration behavior.

## Abbreviations

PC: Unstructured protocerebrum
OL: Optic lobe
ME: Medulla
LO: Lobula
LOP: Lobula plate
LA: Lamina
AME: Accessory medulla
AL: Antennal lobe
MB: Mushroom body
CA: Calyx
PED: Pedunculus
ACA: Accessory calyx
SP: Spur
dL: Dorsal lobelet
vL: Ventral lobelet
LX: Lateral complex
LAL: Lateral accessory lobe
BU: Bulb
GA: Gall
CX: Central complex
FB: Fan-shaped body
EB: Ellipsoid body
PB: Protocerebral bridge
NO: Nodulus
AOTU: Anterior optic tubercle
UU: Upper unit
LU: Lower unit
NU: Nodular unit
ONP: Ocellar neuropil
POTU: Posterior optic tubercle
POC: Posterior optic commissure
IT: Isthmus tract
iLALT: Inter-LAL tract
SOG: Suboesophageal ganglion
AOT: Anterior optic tract

## Acknowledgements

The authors are grateful to Ken Green, David Dreyer and Kristina Brauburger for help with moth collection and maintenance, and to Eva Landgren for assistance with histology. We thank Dr. Erich Buchner, University of Würzburg for providing the anti-Synapsin antibody. We also thank the staff of the HPC cluster MaRC2 at the University of Marburg for their support. This work was funded by the US Airforce Office of Scientific Research (grant number FA9550-14-1-0242) to SH and EW, and the European Research Council (ERC) under the European Union’s Horizon 2020 research and innovation program (grant agreement no. 714599 to S.H., and grant agreement no. 741298 to E.W.).

## References

Amador-Vargas S, Gronenberg W, Wcislo WT, Mueller U. 2015. Specialization and group size: Brain and behavioral correlates of colony size in ants lacking morphological castes. Proc R Soc B Biol Sci 282.

Aso Y, Sitaraman D, Ichinose T, Kaun KR, Vogt K, Belliart-Guérin G, Plaçais PY, Robie AA, Yamagata N, Schnaitmann C, Rowell WJ, Johnston RM, Ngo TTB, Chen N, Korff W, Nitabach MN, Heberlein U, Preat T, Branson KM, Tanimoto H, Rubin GM. 2014. Mushroom body output neurons encode valence and guide memory-based action selection in Drosophila. Elife 3:e04580.

Berg BG, Almaas TJ, Bjaalie JG, Mustaparta H. 1998. The macroglomerular complex of the antennal lobe in the tobacco budworm moth Heliothis virescens: Specified subdivision in four compartments according to information about biologically significant compounds. J Comp Physiol - A Sensory, Neural, Behav Physiol 183:669–682.

Berg BG, Galizia CG, Brandt R, Mustaparta H. 2002. Digital atlases of the antennal lobe in two species of tobacco budworm moths, the oriental Helicoverpa assulta (male) and the American Heliothis virescens (male and female). J Comp Neurol 446:123–134.

Clarac F, Pearlstein E. 2007. Invertebrate preparations and their contribution to neurobiology in the second half of the 20th century. Brain Res Rev 54:113–161.

Common IFB. 1954. A study of the ecology of the adult Bogong moth, Agrotis infusa (Boisd.) (Lepidoptera: Noctuidae), with special reference to its behaviour during migration and aestivation. Aust J Zool 2:223–263.

Cronin TW, Warrant EJ, Greiner B. 2006. Celestial polarization patterns during twilight. Appl Opt 45:5582–5589.

Dreyer D, Frost B, Mouritsen H, Günther A, Green K, Whitehouse M, Johnsen S, Heinze S, Warrant E. 2018. The Earth’s Magnetic Field and Visual Landmarks Steer Migratory Flight Behavior in the Nocturnal Australian Bogong Moth. Curr Biol 28:1–7.

Eaton JL, Tignor KR, Holtzman GI. 1983. Role of moth ocelli in timing flight initiation at dusk. Physiol Entomol 8:371–375.

Green J, Adachi A, Shah KK, Hirokawa JD, Magani PS, Maimon G. 2017. A neural circuit architecture for angular integration in Drosophila. Nature 546:101–106.

Gronenberg W. 2001. Subdivisions of hymenopteran mushroom body calyces by their afferent supply. J Comp Neurol 435:474–489.

Guerra PA, Gegear RJ, Reppert SM. 2014. A magnetic compass aids monarch butterfly migration. Nat Commun 5.

von Hadeln J, Althaus V, Häger L, Homberg U. 2018. Anatomical organization of the cerebrum of the desert locust Schistocerca gregaria. Cell Tissue Res 374:39–62.

Hansson BS, Anton S, Christensen TA. 1994. Structure and function of antennal lobe neurons in the male turnip moth, Agrotis segetum (Lepidoptera: Noctuidae). J Comp Physiol A 175:547–562.

Heinze S. 2014. Polarized Light and Polarization Vision in Animal Sciences. In: Horváth G, editor. Polarized Light and Polarization Vision in Animal Sciences. Springer-Verlag Berlin Heidelberg. p 61–111.

Heinze S, Florman J, Asokaraj S, el Jundi B, Reppert SM. 2013. Anatomical basis of sun compass navigation II: The neuronal composition of the central complex of the monarch butterfly. J Comp Neurol 521:267–298.

Heinze S, Homberg U. 2008. Neuroarchitecture of the central complex of the desert locust: Intrinsic and columnar neurons. J Comp Neurol 511:454–478.

Heinze S, Reppert SM. 2011. Sun compass integration of skylight cues in migratory monarch butterflies. Neuron 69:345–358.

Heinze S, Reppert SM. 2012. Anatomical basis of sun compass navigation I: The general layout of the monarch butterfly brain. J Comp Neurol 520:1599–1628.

Homberg U. 2015. Sky Compass Orientation in Desert Locusts — Evidence from Field and Laboratory Studies. Front Behav Neurosci 9.

Homberg U, Hofer S, Pfeiffer K, Gebhardt S. 2003. Organization and Neural Connections of the Anterior Optic Tubercle in the Brain of the Locust, Schistocerca gregaria. J Comp Neurol 462:415–430.

Homberg U, Montague RA, Hildebrand JG. 1988. Anatomy of antenno-cerebral pathways in the brain of the sphinx moth Manduca sexta. Cell Tissue Res 254:255–281.

Honkanen A, Adden A, da Silva Freitas J, Heinze S. 2019. The insect central complex and the neural basis of navigational strategies. J Exp Biol 222.

Ian E, Berg A, Lillevoll SC, Berg BG. 2016. Antennal-lobe tracts in the noctuid moth, Heliothis virescens: new anatomical findings. Cell Tissue Res 366:23–35.

Immonen EV, Dacke M, Heinze S, el Jundi B. 2017. Anatomical organization of the brain of a diurnal and a nocturnal dung beetle. J Comp Neurol 525:1879–1908.

Ito K, Shinomiya K, Ito M, Armstrong JD, Boyan G, Hartenstein V, Harzsch S, Heisenberg M, Homberg U, Jenett A, Keshishian H, Restifo LL, Rössler W, Simpson JH, Strausfeld NJ, Strauss R, Vosshall LB. 2014. A systematic nomenclature for the insect brain. Neuron 81:755–65.

Iwano M, Hill ES, Mori A, Mishima T, Mishima T, Ito K, Kanzaki R. 2010. Neurons Associated With the Flip-Flop Activity in the Lateral Accessory Lobe and Ventral Protocerebrum of the Silkworm Moth Brain. J Comp Neurol 518:366–388.

Jarriault D, Gadenne C, Lucas P, Rospars JP, Anton S. 2010. Transformation of the sex pheromone signal in the noctuid moth Agrotis ipsilon: From peripheral input to antennal lobe output. Chem Senses 35:705–715.

Jefferis GSXE, Potter CJ, Chan AM, Marin EC, Rohlfing T, Maurer CR, Luo L. 2007. Comprehensive Maps of Drosophila Higher Olfactory Centers: Spatially Segregated Fruit and Pheromone Representation. Cell 128:1187–1203.

Jenett A, Rubin GM, Ngo TTB, Shepherd D, Murphy C, Dionne H, Pfeiffer BD, Cavallaro A, Hall D, Jeter J, Iyer N, Fetter D, Hausenfluck JH, Peng H, Trautman ET, Svirskas RR, Myers EW, Iwinski ZR, Aso Y, DePasquale GM, Enos A, Hulamm P, Lam SCB, Li HH, Laverty TR, Long F, Qu L, Murphy SD, Rokicki K, Safford T, Shaw K, Simpson JH, Sowell A, Tae S, Yu Y, Zugates CT. 2012. A GAL4-Driver Line Resource for Drosophila Neurobiology. Cell Rep 2:991–1001.

el Jundi B, Baird E, Byrne MJ, Dacke M. 2019. The brain behind straight-line orientation in dung beetles. J Exp:1–7.

el Jundi B, Huetteroth W, Kurylas AE, Schachtner J. 2009. Anisometric brain dimorphism revisited: Implementation of a volumetric 3D standard brain in Manduca sexta. J Comp Neurol 517:210–225.

King JR, Christensen TA, Hildebrand JG. 2000. Response characteristics of an identified, sexually dimorphic olfactory glomerulus. J Neurosci 20:2391–2399.

Kinoshita M, Shimohigasshi M, Tominaga Y, Arikawa K, Homberg U. 2015. Topographically distinct visual and olfactory inputs to the mushroom body in the swallowtail butterfly, papilio xuthus. J Comp Neurol 523:162–182.

Klagges BRE, Heimbeck G, Godenschwege TA, Hofbauer A, Pflugfelder GO, Reifegerste R, Reisch D, Schaupp M, Buchner S, Buchner E. 1996. Invertebrate Synapsins: A Single Gene Codes for Several Isoforms in Drosophila. J Neurosci 16:3154–3165.

Kvello P, Løfaldli BB, Rybak J, Menzel R, Mustaparta H. 2009. Digital, three-dimensional average shaped atlas of the Heliothis virescens brain with integrated gustatory and olfactory neurons. Front Syst Neurosci 3:1–14.

Lin C, Strausfeld NJ. 2012. Visual inputs to the mushroom body calyces of the whirligig beetle Dineutus sublineatus: Modality switching in an insect. J Comp Neurol 520:2562– 2574.

Menzel R. 2014. The insect mushroom body, an experience-dependent recoding device. J Physiol Paris 108:84–95.

Molina Y, O’Donnell S. 2008. Age, sex, and dominance-related mushroom body plasticity in the paperwasp Mischocyttarus mastigophorus. Dev Neurobiol 68:950–959.

Montgomery SH, Merrill RM, Ott SR. 2016. Brain composition in Heliconius butterflies, posteclosion growth and experience-dependent neuropil plasticity. J Comp Neurol 524:1747–1769.

Montgomery SH, Ott SR. 2015. Brain composition in Godyris zavaleta, a diurnal butterfly, Reflects an increased reliance on olfactory information. J Comp Neurol 523:869–891.

Mota T, Yamagata N, Giurfa M, Gronenberg W, Sandoz J-C. 2011. Neural Organization and Visual Processing in the Anterior Optic Tubercle of the Honeybee Brain. J Neurosci 31:11443–11456.

Mouritsen H, Derbyshire R, Stalleicken J, Mouritsen OØ, Frost BJ, Norris DR. 2013. An experimental displacement and over 50 years of tag-recoveries show that monarch butterflies are not true navigators. PNAS 110:7348–7353.

Namiki S, Kanzaki R. 2016. The neurobiological basis of orientation in insects: insights from the silkmoth mating dance. Curr Opin insect Sci in press:16–26.

Omoto JJ, Keleş MF, Nguyen BCM, Bolanos C, Lovick JK, Frye MA, Hartenstein V. 2017. Visual Input to the Drosophila Central Complex by Developmentally and Functionally Distinct Neuronal Populations. Curr Biol 27:1098–1110.

Ott SR. 2008. Confocal microscopy in large insect brains: Zinc-formaldehyde fixation improves synapsin immunostaining and preservation of morphology in whole-mounts. J Neurosci Methods 172:220–230.

Paulk AC, Gronenberg W. 2008. Higher order visual input to the mushroom bodies in the bee, Bombus impatiens. Arthropod Struct Dev 37:443–458.

Pfeiffer K, Homberg U. 2007. Coding of Azimuthal Directions via Time-Compensated Combination of Celestial Compass Cues. Curr Biol 17:960–965.

Pfeiffer K, Kinoshita M, Homberg U. 2005. Polarization-sensitive and light-sensitive neurons in two parallel pathways passing through the anterior optic tubercle in the locust brain. J Neurophysiol 94:3903–3915.

Preibisch S, Saalfeld S, Tomancak P. 2009. Globally optimal stitching of tiled 3D microscopic image acquisitions. Bioinformatics 25:1463–1465.

Reppert SM, Guerra PA, Merlin C. 2016. Neurobiology of Monarch Butterfly Migration. Annu Rev Entomol 61:25–42.

Ritz T, Adem S, Schulten K. 2000. A Model for Photoreceptor-Based Magnetoreception in Birds. Biophys J 78:707–718.

Rohlfing T, Brandt R, Maurer CR, Menzel R. 2001. Bee brains, B-splines and computational democracy: Generating an average shape atlas. Proc Work Math Methods Biomed Image Anal:187–194.

Rohlfing T, Maurer CR. 2007. Shape-Based Averaging. IEEE Trans Image Process 16:153– 161.

Schindelin J, Arganda-Carreras I, Frise E, Kaynig V, Longair M, Pietzsch T, Preibisch S, Rueden C, Saalfeld S, Schmid B, Tinevez JY, White DJ, Hartenstein V, Eliceiri K, Tomancak P, Cardona A. 2012. Fiji: An open-source platform for biological-image analysis. Nat Methods 9:676–682.

Schulten K, Swenberg CE, Weiler A. 1978. A Biomagnetic Sensory Mechanism Based on Magnetic Field Modulated Coherent Electron Spin Motion. Zeitschrift fur Phys Chemie 111:1–5.

Schultzhaus JN, Saleem S, Iftikhar H, Carney GE. 2017. The role of the Drosophila lateral horn in olfactory information processing and behavioral response. J Insect Physiol 98:29–37.

Seelig JD, Jayaraman V. 2013. Feature detection and orientation tuning in the Drosophila central complex. Nature 503:262.

Shiozaki HM, Kazama H. 2017. Parallel encoding of recent visual experience and self-motion during navigation in Drosophila. Nat Neurosci 20:1395–1403.

Sinakevitch I, Sjöholm M, Hansson BS, Strausfeld NJ. 2008. Global and local modulatory supply to the mushroom bodies of the moth Spodoptera littoralis. Arthropod Struct Dev 37:260–272.

Sjöholm M, Sinakevitch I, Ignell R, Strausfeld NJ, Hansson BS. 2005. Organization of Kenyon cells in subdivisions of the mushroom bodies of a lepidopteran insect. J Comp Neurol 491:290–304.

Stange G. 1981. The ocellar component of flight equilibrium control in dragonflies. J Comp Physiol □ A 141:335–347.

Stöckl A, Heinze S, Charalabidis A, el Jundi B, Warrant E, Kelber A. 2016. Differential investment in visual and olfactory brain areas reflects behavioural choices in hawk moths. Sci Rep:1–10.

Stöckl AL, Heinze S. 2015. A clearer view of the insect brain — combining bleaching with standard whole-mount immunocytochemistry allows confocal imaging of pigment-covered brain areas for 3D reconstruction. Front Neuroanat 9:1–8.

Stone T, Webb B, Adden A, Weddig N Ben, Honkanen A, Templin R, Wcislo W, Scimeca L, Warrant E, Heinze S. 2017. An Anatomically Constrained Model for Path Integration in the Bee Brain. Curr Biol 27:3069–3085.e11.

Strausfeld NJ. 2012. Arthropod Brains. Cambridge, Massachusetts, London, England: Harvard University Press.

Strausfeld NJ, Hansen L, Li Y, Gomez RS, Ito K. 1998. Evolution, discovery, and interpretations of arthropod mushroom bodies. Learn Mem 5:11–37.

Strube-Bloss MF, Nawrot MP, Menzel R. 2011. Mushroom body output neurons encode odor-reward associations. J Neurosci 31:3129–3140.

Strutz A, Soelter J, Baschwitz A, Farhan A, Grabe V, Rybak J, Knaden M, Schmuker M, Hansson BS, Sachse S. 2014. Decoding odor quality and intensity in the Drosophila brain. Elife 3:e04147.

Turner-Evans D, Wegener S, Rouault H, Franconville R, Wolff T, Seelig JD, Druckmann S, Jayaraman V. 2017. Angular velocity integration in a fly heading circuit. Elife 6:e23496.

Varga AG, Ritzmann RE. 2016. Cellular Basis of Head Direction and Contextual Cues in the Insect Brain. Curr Biol 26:1816–1828.

de Vries L, Pfeiffer K, Trebels B, Adden AK, Green K, Warrant E, Heinze S. 2017. Comparison of Navigation-Related Brain Regions in Migratory versus Non-Migratory Noctuid Moths. Front Behav Neurosci 11:1–19.

Warrant E, Dacke M. 2016. Visual Navigation in Nocturnal Insects. Physiology 31:182–192.

Warrant E, Frost B, Green K, Mouritsen H, Dreyer D, Adden A, Brauburger K, Heinze S. 2016. The australian bogong moth Agrotis infusa: A long-distance nocturnal navigator. Front Behav Neurosci 10.

Williams JLD. 1975. Anatomical studies of the insect central nervous system: A ground-plan of the midbrain and an introduction to the central complex in the locust, Schistocerca gregaria (Orthoptera). J Zool 176:67–86.

Wolff T, Iyer NA, Rubin GM. 2015. Neuroarchitecture and neuroanatomy of the Drosophila central complex: A GAL4-based dissection of protocerebral bridge neurons and circuits. J Comp Neurol 523:997–1037.

Wunderer H, de Kramer JJ. 1989. Dorsal ocelli and light-induced diurnal activity patterns in the arctiid moth Creatonotos transiens. J Insect Physiol 35:87–95.

Zeil J, Ribi WA, Narendra A. 2014. Polarisation Vision in Ants, Bees and Wasps. In: Horváth G, editor. Polarized Light and Polarization Vision in Animal Sciences. Berlin, Heidelberg: Springer. p 41–60.

Zeller M, Held M, Bender J, Berz A, Heinloth T, Hellfritz T, Pfeiffer K. 2015. Transmedulla Neurons in the Sky Compass Network of the Honeybee (Apis mellifera) Are a Possible Site of Circadian Input. PLoS One 10:1–25.

Zhao XC, Chen QY, Guo P, Xie GY, Tang QB, Guo XR, Berg BG. 2016. Glomerular identification in the antennal lobe of the male moth Helicoverpa armigera. J Comp Neurol 524:2993–3013.

Zorovic M, Hedwig B. 2011. Processing of species-specific auditory patterns in the cricket brain by ascending, local, and descending neurons during standing and walking. J Neurophysiol 105:2181–2194.

Insect Brain Database: https://hdl.handle.net/20.500.12158/SIN-0000002.1

